# A hierarchical Bayesian interaction model to estimate cell-type-specific methylation quantitative trait loci incorporating priors from cell-sorted bisulfite sequencing data

**DOI:** 10.1101/2024.02.01.578272

**Authors:** Youshu Cheng, Biao Cai, Hongyu Li, Xinyu Zhang, Gypsyamber D’Souza, Sadeep Shrestha, Andrew Edmonds, Jacquelyn Meyers, Margaret Fischl, Seble Kassaye, Kathryn Anastos, Mardge Cohen, Bradley E Aouizerat, Ke Xu, Hongyu Zhao

**Author notes:** **Correspondence:** Ke Xu, MD, PhD, Professor of Psychiatry, Yale School of Medicine, Hongyu Zhao, PhD, Ira V. Hiscock Professor of Biostatistics, of Genetics, and of Statistics and Data Science, Department of Biostatistics, Yale School of Public Health. Electronic address.

## Abstract

**Background:** Methylation Quantitative Trait Loci (meQTLs) are chromosomal regions that harbor genetic variants affecting DNA methylation levels. The identification of meQTLs can be accomplished through quantifying the effects of single nucleotide polymorphisms (SNPs) on DNA methylation levels, and these inferred meQTLs can shed light on the complex interplay between the genome and methylome. However, most meQTL studies to date utilize bulk methylation datasets composed of different cell types that may have distinct methylation patterns in each cell type. Current technological challenges hinder the comprehensive collection of large-scale, cell-type-specific (CTS) methylation data, which limits our understanding of CTS methylation regulation. To address this challenge, we propose a hierarchical Bayesian interaction model (HBI) to infer CTS meQTLs from bulk methylation data.

**Results:** Our HBI method integrates bulk methylations data from a large number of samples and CTS methylation data from a small number of samples to estimate CTS meQTLs. Through simulations, we show that HBI improves the estimation (accuracy and power) of CTS genetic effects on DNA methylation. To systematically characterize genome-wide SNP-methylation level associations in multiple cell types, we apply HBI to bulk methylation data measured in peripheral blood mononuclear cells (PBMC) from a cohort of 431 individuals together with flow-sorted cell-derived methylation sequencing (MC-seq) data measured in isolated white blood cells (CD4+ T-cells, CD8+ T-cells, CD16+ monocytes) for 47 individuals. We demonstrate that HBI can identify CTS meQTLs and improve the functional annotation of SNPs.

**Conclusions:** HBI can incorporate strong and robust signals from MC-seq data to improve the estimation of CTS meQTLs. Applying HBI to link the methylome and genome data helps to identify biologically relevant cell types for complex traits.

## Background

DNA methylation (DNAm) is one of the most widely studied epigenetic modifications that capture the cumulative effects of environmental and genetic factors. DNAm regulates cellular differentiation and gene expression and plays a key role in human development and disease etiology [1, 2]. Single nucleotide polymorphisms (SNPs) associated with DNAm levels are known as methylation quantitative trait loci (meQTLs) [3-6], which capture and represent the complex interplay between the genome and methylome.

To reveal cellular mechanisms for DNAm patterns and their link to complex traits, it is important to study cell-type-specific (CTS) genetic effects on DNAm (CTS-meQTL). For example, SNP rs174548, which is mapped on *FADS1*, a key enzyme in the metabolism of fatty acids, is associated with asthma [1]. At the same time, its effect on methylation at cg21709803 is the strongest in CD8+ T-cells. These results suggest a possible effect of rs174548 on asthma via immune dysregulation and fatty acid metabolism through methylation in CD8+ T-cells [1]. However, most meQTL studies to date have used bulk samples composed of distinct cell types [7-9]. MeQTLs identified from bulk DNAm samples reflect the aggregated genetic effects across all cell types, which provide no insights for genetic regulations in individual cell types. This approach is especially problematic for rare or less abundant cell types. The high cost and technical limitations for both cell sorting and single-cell DNAm approaches hinder the collection of large-scale, CTS methylation profiles, and limit our ability to move meQTL studies from the “bulk level” to the “cell type level.”

Given the difficulty in generating large-scale CTS methylome data to directly estimate CTS effects and the broad availability of many bulk methylation datasets, several statistical methods have been developed to infer CTS meQTLs from bulk data. These methods can be classified into two categories. Methods in the first category estimate sample-level CTS DNAm profiles from bulk data in the first step, and then test the associations with outcomes of interest using the deconvoluted data for each cell type. Tensor Composition Analysis (TCA), a frequentist approach in this category, was originally designed to identify CTS differentially methylated CpG sites in epigenome-wide association studies of phenotypes (CTS-EWAS) [10]. There is also a similar algorithm designed for gene expression data [11], named Bayesian MIND (bMIND), which further incorporates information from single-cell RNA sequencing (scRNA-seq) data as a prior to refine the estimation of CTS expression for each bulk sample. bMIND innovatively integrates large-scale bulk data and small-scale CTS expression data from scRNA-seq to estimate CTS expression for large-scale bulk samples. In contrast, methods in the second category are based on an interaction model to test the interaction between cell type fractions and variables of interest without deconvolution. Examples include CellDMC [12], which focuses on the interaction between cell type fractions and phenotypes (CTS-EWAS). Westra et al. also proposed an interaction model to estimate CTS expression quantitative trait loci (CTS-eQTL) [13].

Here we introduce a hierarchical Bayesian interaction model (HBI) to infer CTS meQTLs from bulk methylation data. Our model allows the incorporation of cell-type specific DNAm data from a relatively small number of samples to improve the performance of HBI. Compared with bMIND, which utilizes Bayesian techniques to infer the posterior mean of sample-level CTS expression (or as easily for methylation), the goal of HBI is instead to infer the posterior mean of CTS genetic effects by placing sparse hierarchical priors on regression coefficients for the interaction terms. In our model, we employ hierarchical double-exponential priors to induce different shrinkage for different variables, which corresponds to the Bayesian adaptive lasso [14]. If cell-type-specific data are available for a small number of samples (e.g., 5%-10% of the sample size in bulk data), the algorithm can incorporate this information to further refine the estimates for CTS genetic effects in the larger-scale bulk samples. In our case, cell-sorted sequencing data is used to derive CTS DNAm and since it offers the unique advantage of directly measuring CTS methylomes, incorporating strong and robust signals from the MC-seq data will improve the estimation of CTS-meQTLs.

We show in simulations that HBI improves the estimation of CTS genetic effects when compared to other state-of-the-art methods [10-12]. We apply our method to identify cis CTS-meQTL using data from samples in the Women’s Interagency HIV Study (WIHS) (n_bulk_=431, n_CTS_=47) [15]. To demonstrate the utility of our method, we use an independent meQTL dataset derived from CTS methylation data [1] to evaluate the replication of HBI-identified signals. Finally, we perform downstream analyses to improve the annotation of functional genetic variants and to reveal the cellular specificity of complex traits.

## Results

### Estimation of CTS-meQTLs using HBI

A linear regression framework including interaction terms between genotype/phenotype and estimated cell type fractions has been applied to identify CTS-QTL or CTS-differentially methylated CpGs [12, 13]. Here, based on this idea, we propose HBI to incorporate prior information from CTS DNAm data and to improve the estimation of CTS-meQTLs (**Figure 1**). We place hierarchical double-exponential priors on regression coefficients for the interaction terms:

**Figure 1:**
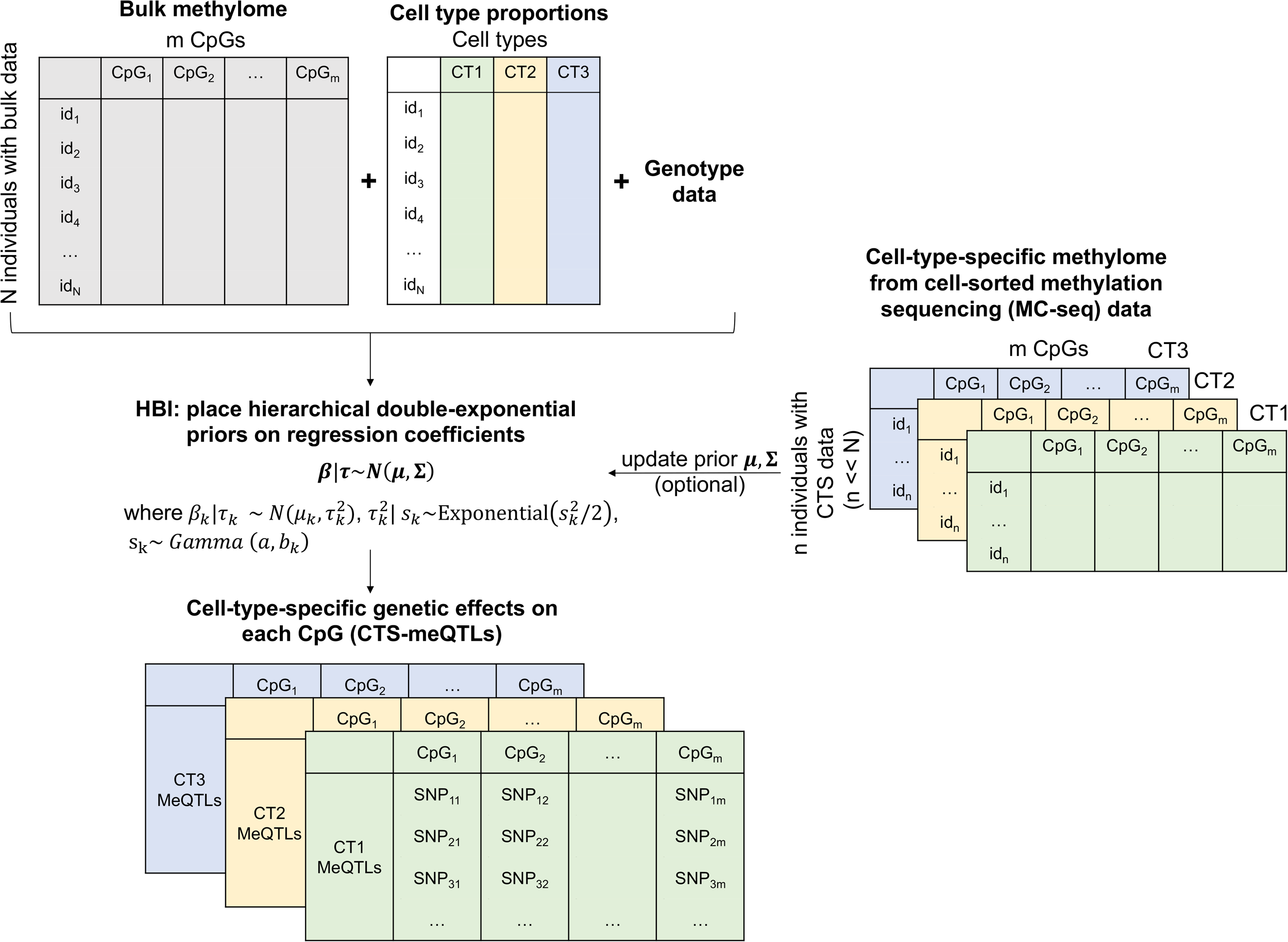
Overview of the hierarchical Bayesian interaction model (HBI) to infer cell type specific (CTS) meQTLs. With bulk methylation data and cell type proportions (we present an example of three cell types: CT1, CT2, CT3), HBI employs an interaction model with sparse hierarchical priors placed on the regression coefficients for the interaction terms. If the CTS DNA methylation data (in our case, generated by methylation capture-sequencing using cells sorted from PBMC using flow cytometry) are available for a small group of samples, HBI will further incorporate the information into priors to refine the estimates for CTS genetic effects in the larger-scale bulk samples.

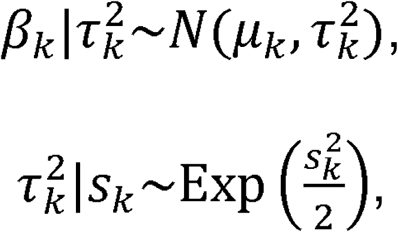

where *β*_*k*_ is the regression coefficient on the interaction term between genotype and cell type proportion for the *k* th cell type. Marginalizing over 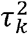, *β*_*k*_ conditional on *s*_*k*_ follows a double-exponential distribution:

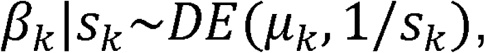

where parameter *s*_*k*_ controls the degree of shrinkage. If *s*_*k*_ is a fixed value for *k* = 1,2,…,*K*, each variable will be shrunk to the same degree. Here we model *s*_*k*_ as a hyperparameter to allow for variable-specific penalty:

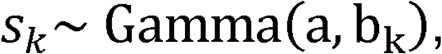

Where *a* and *b*_*k*_ are chosen based on empirical experiments (**Methods**).

In the case where only bulk data are available, we set µ_*k*_*=* 0 for *k* =1,2,…*K* and the model would be similar to the adaptive Lasso approach [14, 16]: the regression coefficients for interaction terms are shrunk to 0 and the degree of shrinkage differs for different variables. Such shrinkage helps to take the sparsity of genetic effects into consideration. When the CTS methylomes are available for a small number of samples, we can first get a rough estimate of the genetic effect in the *k* th cell type using the small set. Then instead of setting *µ*_*k*_ = 0 and shrinking the coefficient to zero, we can shrink it to a more meaningful value by updating the prior mean *µ*_k:_

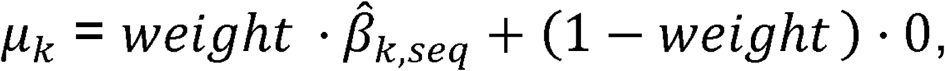

where *µ*_*k*_ is a weighted sum between 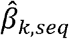 (observed results from CTS methylomes) and zero (prior beliefs), while *weight* = 1−*p*_*adjust*_ and *p*_*adjust*_ is the p-value adjusted using the Bonferroni correction (**Methods**). Similar to other studies that propose weights based on posterior probabilities [17], the weights in our model are assigned based on p-values as p-value is a probability measuring the evidence against the null hypothesis (*β*_*k,seq*_ = 0) [18] and can reflect the stability of the estimator 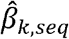. Intuitively, a small p-value closer to zero indicates 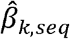 estimated using the CTS DNAm data is relatively strong. In this case, we shrink the coefficient more towards 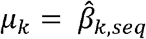. In contrast, a large p-value closer to one indicates 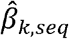 is not significantly different from zero, and thus we shrink it more towards *µ*_*k*_ = 0.

Along with updating prior means, we can also update prior variances:

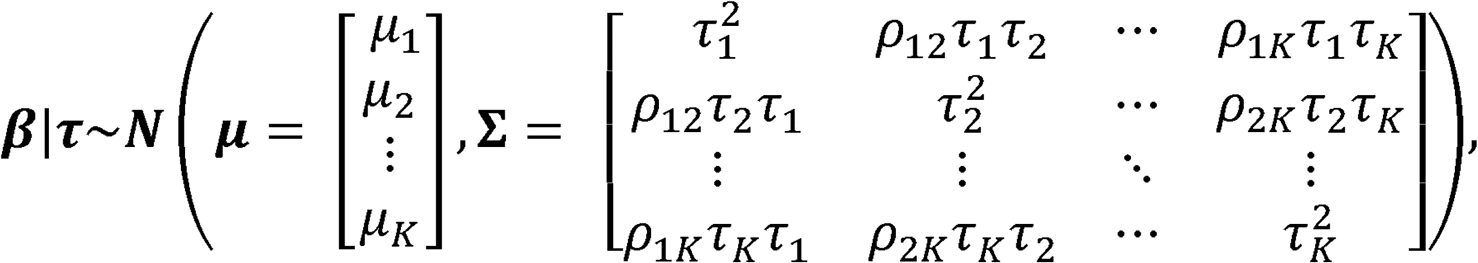

Where *ρ*_*jk*_ can be updated as the genetic correlation between cell type *k* methylation and cell type *j* methylation, which can also be estimated from the CTS methylomes provided. The prior variance without CTS data can be seen as a special case with all *ρ*_*jk*_ *=* 0. Of note, as the detection of genetic effects is always much harder in less abundant cell types, the incorporation of the estimated genetic correlation aims to improve the power in the less abundant cell types by borrowing information from the more abundant cell types.

More details of our model are summarized in **Methods**. We note the following key features for HBI: (1) because only a few SNPs among hundreds of SNPs surrounding a CpG may have detectable effects, placing a sharp prior centered at 0 helps to take the sparsity of genetic effects into consideration; (2) local shrinkage parameters *s*_*k*_ and 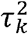 make the algorithm more flexible: the degree of shrinkage could differ among variables; and (3) priors could be updated to incorporate information from CTS DNAm data, if they are available for a small group of samples.

### HBI improves performance in simulations

We evaluated the performance of HBI in estimating CTS-meQTLs through extensive simulations. We considered three scenarios: (1) there are genetic effects only in the major/most abundant cell type; (2) there are genetic effects only in the minor/least abundant cell type; and (3) there are correlated genetic effects in all cell types. We compared HBI to other state-of-the-art methods: bMIND, TCA, and the basic interaction model fitted by ordinary least squares (OLS) (**Methods**). In each scenario, we assessed the correlation between the estimated and true effect sizes, mean squared error (MSE) between the estimated and true effect sizes, power, and false discovery rate (FDR) as a function of the proportion of causal SNPs.

HBI improved the point estimation of CTS-meQTLs by achieving higher correlation and lower MSE (**Figure 2**). For example, in scenario 1 when the proportion of causal SNPs fixed at 10%, the median of correlation across 10 simulations was 0.72 for bMIND with only bulk data (denoted as “bMIND”), 0.71 for bMIND with CTS data incorporated (denoted as “bMIND_CTS-prior”), 0.68 for TCA, 0.77 for basic interaction model, 0.94 for HBI with only bulk data (denoted as “HBI”), and 0.94 for HBI with CTS data incorporated (denoted as “HBI_CTS-prior”). Across all scenarios, HBI generally achieved higher power compared with other methods. We note that in scenario 1, when genetic effects only occur in the most abundant cell type, further incorporating CTS DNAm data to update priors had comparable power to the case without incorporating CTS DNAm data. For example, in scenario 1 when the proportion of causal SNPs fixed at 10%, the median of power across 10 simulations was both 0.65 for HBI with only bulk data (denoted as “HBI”) and HBI with CTS data incorporated (denoted as “HBI_CTS-prior”). In contrast, in scenarios 2 and 3, when there were genetic effects in the minor/least abundant cell type, incorporating information from CTS DNAm data helped to improve the power. For example, in scenario 2 when the proportion of causal SNPs fixed at 10%, the median of power across 10 simulations was 0.15 for HBI with only bulk data (denoted as “HBI”), and 0.24 for HBI with CTS data incorporated (denoted as “HBI_CTS-prior”). In each scenario, we varied the proportion of causal SNPs from 10% to 20% to 40%, to compare the performance of these methods when the genetic effects became more polygenic. As expected, the power for all methods decreased as the proportion of causal SNPs increased. When the overall genetic effect (heritability) was fixed and diluted on a larger number of SNPs, it generally became harder to detect signals. We also note that the performance for bMIND and TCA was a result of fitting conditional models (**Methods**). In the case of fitting marginal models for bMIND and TCA, we observed inflated FDR, especially in scenarios 1 and 2 when there were genetic effects only in one single cell type (**Supplementary Figure 1**).

**Figure 2:**
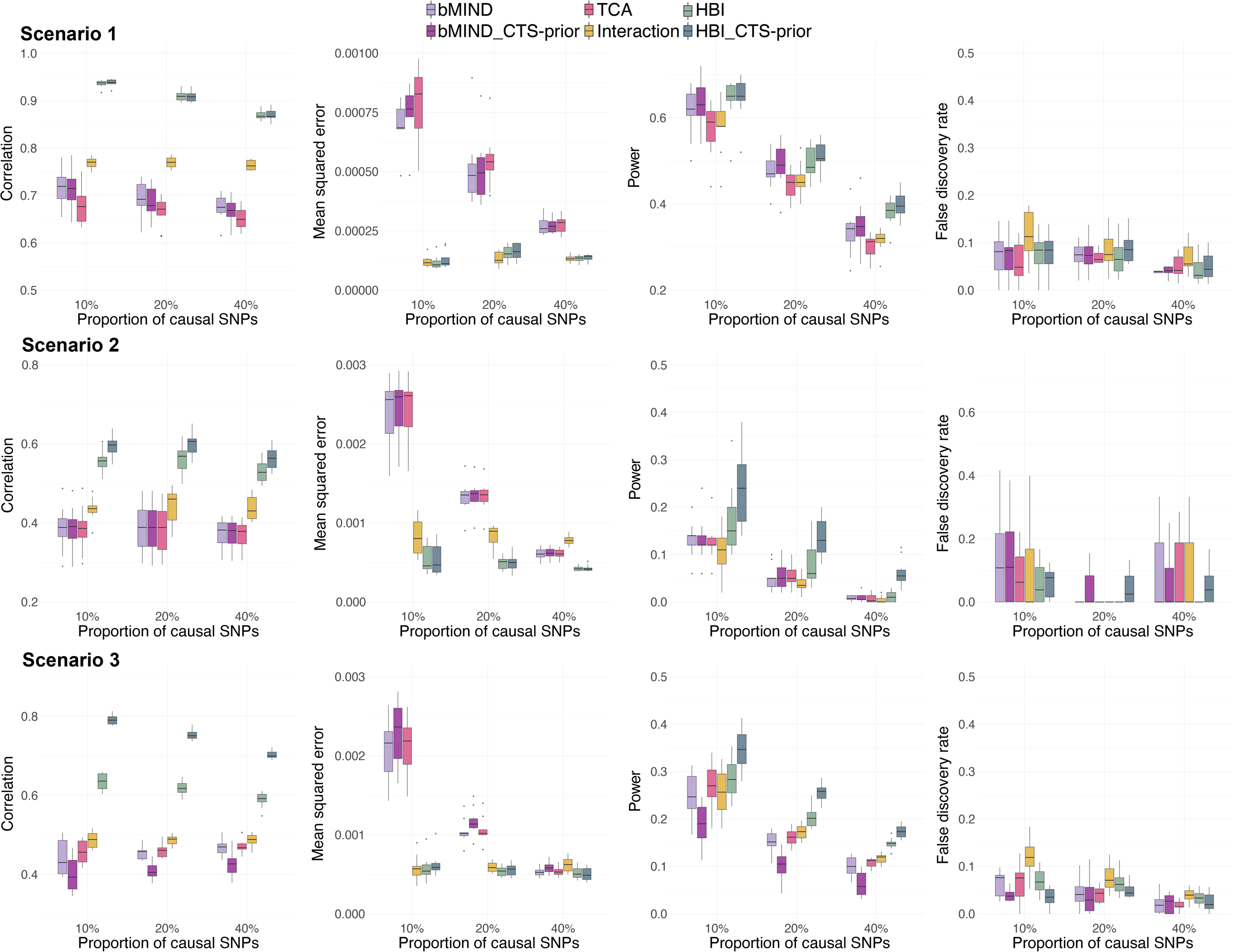
Comparisons of performance in estimating cell type specific (CTS)-meQTLs. From top to bottom: scenarios with genetic effects only in the most abundant cell type (Scenario 1), only in the least abundant cell type (Scenario 2), and with correlated genetic effects in all cell types (Scenario 3) are shown. From left to right: correlation between estimated and true effect sizes, mean squared error (MSE) between estimated and true effect sizes, power, and false discovery rate (FDR) as a function of the proportion of causal SNPs. HBI_CTS-prior, bMIND_CTS-prior represent the version of the corresponding methods with CTS DNA methylation data incorporated.

Of note, all methods included here relied on cell type proportions, but in reality the biological “ground truth” of the cell type proportions is rarely available and the computationally estimated proportions [19, 20] are used directly, which introduces additional noise. Therefore, to further evaluate the robustness of all methods when “noisy” cell type proportions (random error was added to the true proportions) were given, we repeated the simulation scenario 3 but with noisy proportions inputted for all methods (**Methods**). With the increase in the noise added to cell type proportions, the correlation and power decreased while the MSE increased (**Supplementary Figure 2**), as expected. HBI was generally robust in this “noisy” setting by achieving the highest correlation and power and lowest MSE among all the methods considered.

### Genome-wide CTS-meQTLs identification using HBI

To identify genome-wide CTS-meQTLs, we applied HBI to the WIHS cohort with matched genotype data and bulk DNAm data measured in peripheral blood mononuclear cells (PBMC) using the Illumina HumanMethylation EPIC beadchip (n=431) (**Supplementary Table 1**). Furthermore, for a separate group of WIHS participants (n=47), one aliquot of PBMC underwent CTS separation to obtain CD4+ T-cells (n=28), CD8+ T-cells (n=28), or monocytes (n=27). The demographic characteristics of the WIHS participants are described in **Supplemental Table 1**. DNAm from each sorted cell type was profiled using Agilent SureSelectXT Methyl-seq, and the derived priors from these CTS DNAm data were incorporated in HBI (**Methods**). Significant *cis*-meQTLs were selected as those reaching genome-wide significance level (p<6E-12; Bonferroni correction).

HBI identified a total of 122,387 significant meQTLs in CD4+ T-cells, 34,310 in CD8+ T-cells, 25,020 in natural killer cells, 26,972 in B cells, 36,919 in monocytes, and 12,231 in granulocytes (**Figure 3A**) (**Supplementary Table 2**). To replicate our identified CTS-meQTLs, we used publicly available data for meQTLs in isolated white blood cell subsets (CD4+ T-cells, CD8+ T-cells, monocytes) (n=60 individuals) [1], and we defined replicated meQTLs as those with p<0.05 and consistent direction of effect in this replication sample (**Figure 3B**). We showed that among the shared SNP-CpG pairs in the replication sample, 98.2-98.4% had a consistent direction of effect and 79.0–93.9% were replicated (**Figure 3C**). Of note, in all cell types, more than 99% of significant pairs (p<0.05) had a consistent direction of effect (Rep/Sig), indicating a high level of consistency between our results in the WIHS sample and those in the replication sample.

**Figure 3:**
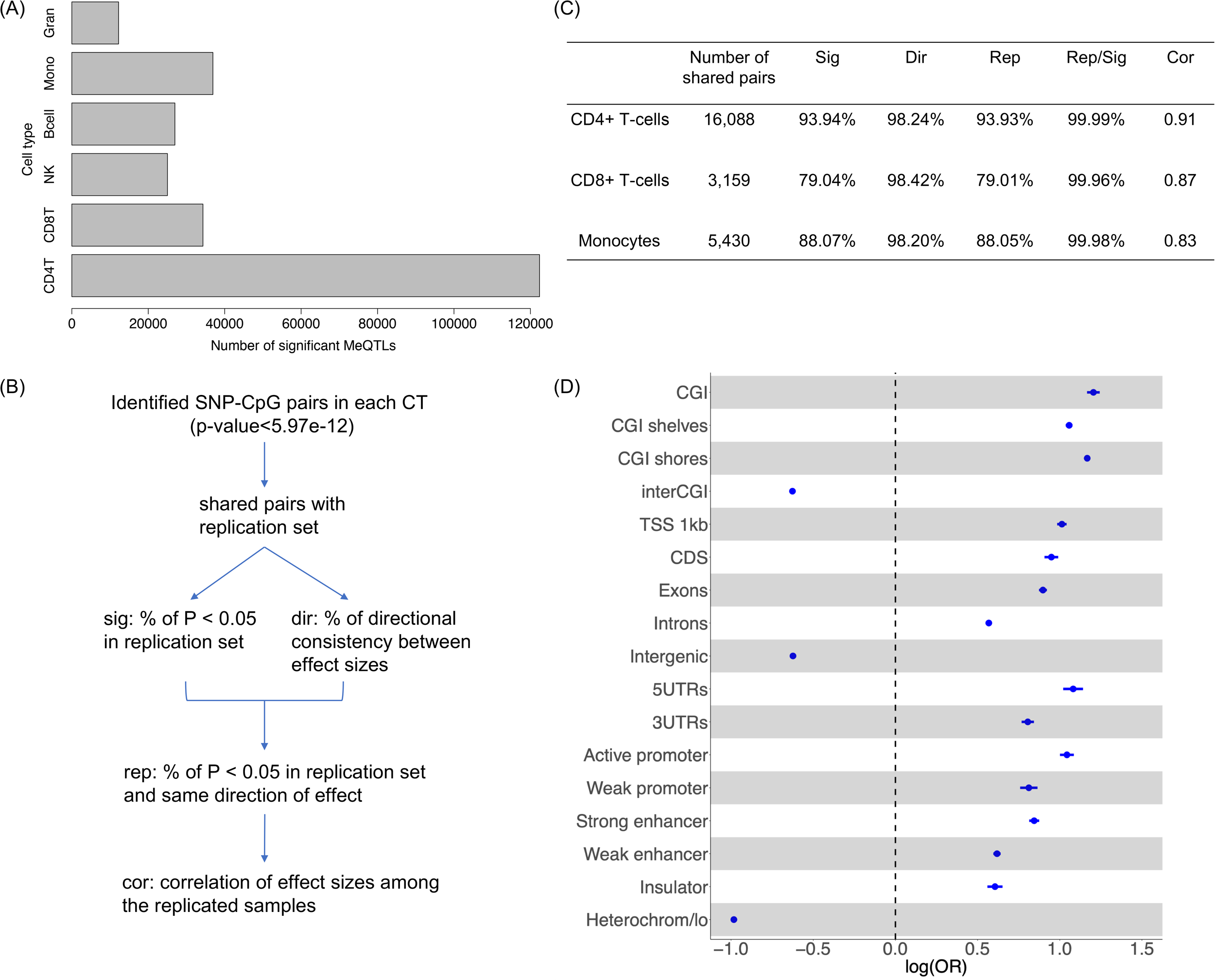
Overview of cell type specific (CTS)-meQTLs identification using the hierarchical Bayesian interaction model (HBI). **(A)** Bar chart shows the number of HBI-identified meQTLs in each cell type (p<6E-12). **(B)** Flow chart indicates the replication of HBI identified CTS-meQTLs in an independent dataset for meQTLs in isolated white blood cell subsets (CD4+ T-cells, CD8+ T-cells, monocytes). **(C)** Table summarizes the replication results. **(D)** Functional enrichment for meQTLs across all cell types in CpG island (CGI) regions, gene body regions and gene regulatory regions. The logarithm of odds ratio (OR) with 95% confidence interval is presented. TSS 1kb: <1kb upstream of the transcription start site (TSS); CDS: coding sequence; UTR: untranslated exon region; Heterochrom/lo: regions that exhibit heterochromatic or heterochromatin-like characteristics; CD4T: CD4+ T-cells; CD8T: CD8+ T-cells; NK: natural killer cells; Mono: monocytes; Gran: granulocytes.

Integrating annotations of CpG islands (CGI), genomic functional regions, and open chromatin states with our derived CTS-meQTL, we observed that compared with SNPs that are not meQTLs (non-meQTLs), our identified meQTLs across all cell types were enriched in important regulatory regions, such as active promotors and strong enhancers (**Figure 3D**). Conversely, our meQTLs were depleted in regions with few active genes, including intergenic regions and regions with heterochromatic characteristics, as previously reported [7]. Of note, meQTLs identified in each cell type exhibited similar functional enrichment patterns and are summarized in **Supplementary Figure 3**.

Using QIAGEN Ingenuity Pathway Analysis (IPA) to perform pathway enrichment analyses of genes mapped by the significant meQTLs in each cell type [21], we found that the antigen presentation pathway was significant in multiple cell types: CD4+ T-cells (p=2.95E-05), CD8+ T-cells (p=1.12E-09), B cells (p=9.55E-06), natural killer cells (p=7.94E-07) and monocytes (p=1.41E-10) (**Supplementary Table 3**). Other identified pathways included the pulmonary fibrosis idiopathic signaling pathway in CD4+ T-cells (p=9.33E-06), the multiple sclerosis signaling pathway and the IL-15 production pathway in CD8+ T-cells (p=9.55E-07 and p=3.63E-05, respectively). These significant pathways indicated that the identified CTS-meQTLs by HBI might play a role in regulating immunity-related functions and activities.

### CTS meQTLs colocalize with risk variants for complex traits

While for most meQTLs the direct impact on complex traits has not been widely reported [22], there have been studies showing that some meQTLs are associated with complex traits and may identify underlying pathways and mechanisms related to diseases [7, 8, 23]. To systematically identify potential associations between meQTLs and complex traits, we applied HyPrColoc (Hypothesis Prioritization for multi-trait Colocalization) [24] to perform an meQTL-GWAS colocalization analysis in each cell type. We integrated the HBI-identified CTS-meQTLs with 57 GWAS datasets in three categories of blood cell counts, cardiometabolic, immune, and allergy [25].

A total of 2,972 significant meQTL-GWAS colocalizations (posterior probability for colocalization (PPFC) > 0.50) were identified across all GWASs and cell types (**Supplementary Table 4A-4F**). Taking a further look into the number of meQTL-GWAS colocalizations per trait across all cell types, we found that GWAS traits in the category of blood cell counts had a larger number of colocalizations compared with traits in other categories (**Supplementary Figure 4**). This abundance of colocalizations was expected as the *cis*-meQTLs were identified in cell types from whole blood. To further illustrate how the meQTL-GWAS colocalizations could differ across cell types, we summarize one example in **Figure 4**. The variant rs2395178 in the *HLA-DRA* gene was identified as a CD8+ T-cell specific meQTL for cg00886432 (p=5.46E-12). As expected, we observed that rs2395178 showed a stronger correlation with DNAm in participants with a high abundance of CD8+ T-cells (**Figure 4A**). Meanwhile, our colocalization analyses revealed that rs2395178 was colocalized between methylation at cg00886432 in CD8+ T-cells and type I diabetes (T1D) (PPFC=0.9802) (**Figure 4B**), while no significant colocalization was observed in other cell types. Of note, polymorphisms at the *HLA-DQ* and *HLA-DR* regions have been recognized as the major genetic determinants of T1D [26]. Taken together, these results suggest that integrating DNA methylome and genome data may help link *HLA-DR* gene function in CD8+ T-cells to T1D.

**Figure 4:**
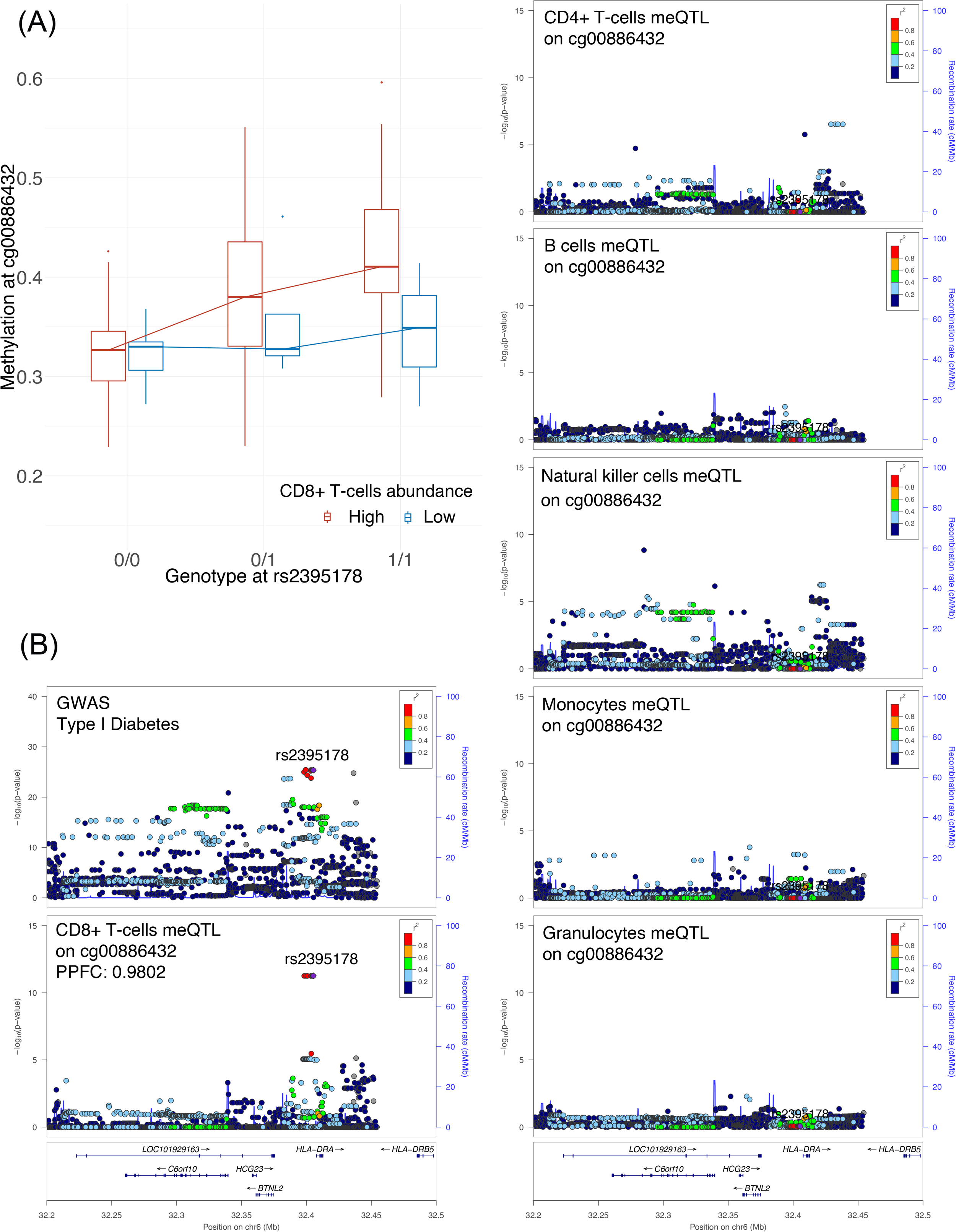
Example of the rs2395178-cg00886432 locus and colocalization results with type I diabetes (T1D). **(A)** An association plot for the rs2395178-cg00886432 locus, separated into individuals with high and low abundance of CD8+ T-cells (above and below the median, respectively). The y axis shows methylation beta-values, while the x axis shows genotypes. **(B)** LocusZoom plots for the association of rs2395178 (mapped to *HLA-DRA*) with phenotypes/molecular traits. Panels illustrate the association of the SNP with GWAS T1D, cg00886432 meQTL signal in CD8+ T-cells, CD4+ T-cells, B cells, natural killer cells, monocytes, and granulocytes. The genetic variant rs2395178 was identified as a colocalized SNP between T1D and cg00886432 meQTL signal in CD8+ T-cells (posterior probability for colocalization (PPFC) is shown).

### MeQTL-GWAS colocalizations exhibit enrichment in trait-relevant cell types

To further investigate meQTL associations with traits in multiple cell types, we performed enrichment analyses to study if the meQTL-GWAS colocalizations for each trait were enriched in certain cell types. Specifically, for each trait we defined the enrichment score in one cell type as the ratio between the percentage of colocalized GWAS-meQTLs in that cell type and the percentage of meQTLs in that cell type (see **Methods**). As the absolute number of colocalizations in each cell type was largely driven by the number of identified meQTLs in that cell type and cannot be compared directly, here the enrichment score was defined as the ratio between two percentages, which allowed us to compare this value across different cell types. We further excluded granulocytes due to the low number of colocalizations identified across traits (**Supplementary Figure 4B**), which indicated the less stable signals identified in this least abundant cell-type. We also evaluated the enrichment score for meQTLs derived at the bulk PBMC level (**Supplementary Table 4G**) to further evaluate whether CTS-meQTLs can reveal more cell-specific information.

We summarize the traits with colocalizations enriched in at least one cell type in **Figure 5A**, which we listed out in **Supplementary Table 5A**. To evaluate whether the enrichment results matched existing biological knowledge, we performed heritability enrichment analyses across the same GWAS traits using GenoSkyline-Plus [27], which could be viewed as an independent tool to identify biologically relevant cell types for complex traits. We found that our significant findings generally agreed with the results of heritability enrichment analyses: 85.2% of our identified cell types with enriched colocalizations were replicated by GenoSkyline-Plus (**Supplementary Table 5B**). This indicates that the colocalizations between HBI-identified CTS-meQTLs and GWASs do help to reveal biologically relevant cell types for complex traits. For example, high-density lipoprotein cholesterol (HDL) had colocalization enrichment in monocytes: monocytes only covered 1.5% of the total meQTLs but accounted for 9.5% of the colocalized meQTLs (enrichment=6.32; p=1.23E-07) (**Figure 5B**). Of note, GenoSkyline-Plus also identified heritability enrichment in monocytes for HDL (p=1.31E-06) (**Supplementary Table 5B**). In T1D, we identified colocalization enrichment in CD8+ T-cells (enrichment=6.15; p=1.95E-05) (**Figure 5B**), while the heritability for this trait was also enriched in CD8+ T-cells (p=4.77E-02) (**Supplementary Table 5B**). This finding is further supported by the evidence that autoreactive CD8+ T-cells play a fundamental role in the progression of T1D by the destruction of pancreatic beta cells [28]. In addition, asthma was enriched in colocalizations derived from CD4+ T-cells (enrichment=4.34; p=5.14E-19) and CD8+ T-cells (enrichment=3.10; p=4.91E-05) meQTLs, which was also replicated by GenoSkyline-Plus (p=3.43E-03 and p=3.92E-05, respectively) (**Supplementary Table 5B**). Interestingly, the association between meQTLs and asthma has been investigated by Hawe et al., who also employed colocalization and reported a shared causal variant rs174548 for methylation at cg21709803 in CD8+ T-cells and asthma [1]. Here our colocalization results in CD8+ T-cells replicated and extended their findings by identifying a nearby risk variant rs174587 (PPFC=0.858) (**Supplementary Table 4B**), which impacts both DNAm at cg21709803 and asthma.

**Figure 5:**
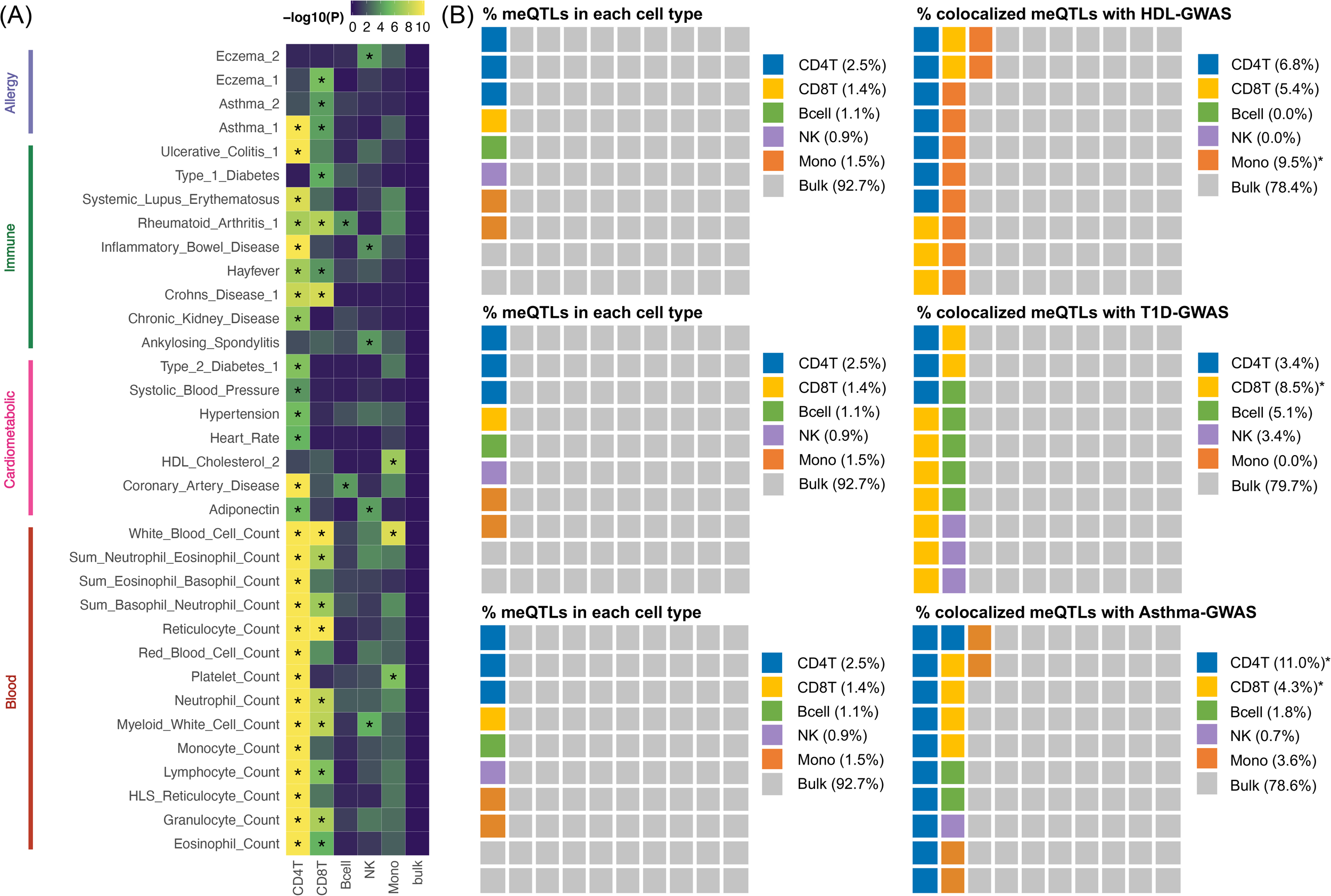
Enrichment analyses for MeQTL-GWAS colocalizations. **(A)** Colocalization enrichment results across six cell types for traits with colocalizations enriched in at least one cell type. Asterisks highlight significance after Bonferroni correction. **(B)** Examples of colocalization enrichments in three traits. From left to right: the percentage of meQTLs covered by each cell type, and the percentage of colocalized meQTLs in that cell type. GWAS: genome-wide association studies; CD4T: CD4+ T-cells; CD8T: CD8+ T-cells; NK: natural killer cells; Mono: monocytes; bulk: a mixture of cell types from peripheral blood mononuclear cells (PBMC).

From **Figure 5B**, we show that more bulk meQTLs were identified than CTS meQTLs, which was consistent with simulations (**Supplementary Figure 5**), but we observed no colocalization enrichments (**Figure 5A**). This suggests that the meQTLs identified in bulk tissue are a mixture of signals from different cell types, thus masking the CTS information. Altogether, those results suggest that our identified CTS meQTLs can provide more insight into the cellular specificity of complex traits and aid the characterization of trait etiology.

## Discussion

We have developed HBI to infer CTS meQTLs from bulk methylation data, with priors derived from CTS methylation data in a small group of samples. As far as we are aware, our model is the first one to integrate large-scale bulk DNAm data and small-scale CTS DNAm data to estimate CTS-meQTLs. We show through simulations that HBI improves the estimation of CTS genetic effects. Applying our method to samples contributed by participants from the WIHS cohort [15], we systematically characterized the genome-wide SNP-CpG associations in multiple cell types of PBMCs. Through colocalization and enrichment analyses, we demonstrate the utility of HBI to improve the annotation of functional genetic variants and enhance the understanding of the cellular specificity of complex traits.

We considered extensive simulation scenarios to compare the performance of different methods in detecting CTS QTLs. As TCA and bMIND were initially developed to detect differentially expressed or differentially methylated signals between comparison groups (e.g., cases versus controls) [10, 11], the differential effect was on a single phenotype of interest in their simulations. In contrast, in our simulations the genetic effects on a CpG were distributed across a number of SNPs and each SNP carried a small effect. Therefore, our simulation aims to detect all causal SNPs, which is more challenging than detecting the association with one single phenotype, and the simulation results may offer a more comprehensive evaluation of the performance of different methods to detect CTS-QTLs than those considered in other studies [10, 11, 29].

The simulation results show that all methods had decreased performance in scenario 2 (the least abundant cell type harbored genetic effects). This is not surprising as the information from rare cell types is in general more limited in a bulk sample, and thus the statistical instability for estimating signals in rare cell types is much larger than that in abundant cell types. In this case, incorporating CTS DNAm information did help to alleviate this problem; we show that HBI with CTS information incorporated into priors (i.e., HBI_CTS-prior) was more powerful than other methods. Specifically, to improve the power to infer meQTL in rare cell types by borrowing information from more abundant cell types, we used the small group with CTS methylation data to estimate *ρ*_*jk*_, the genetic correlation between cell type *k* methylation and cell type *j* methylation, and incorporated the estimated genetic correlation into the prior variance. As cell-sorted MC-seq data offer the unique advantage of directly measuring CTS methylomes, incorporating strong and robust signals from such data improves the estimation of CTS-meQTLs, especially in rare cell types.

We observed inflated FDR when fitting the marginal model for bMIND and TCA. As discussed by the authors of TCA, deconvoluted CTS methylation profiles are expected to be correlated among different cell types [30], which results in false discoveries in the non-causal cell type when using the marginal test model. To mitigate this problem, the TCA authors advised applying a marginal conditional test to account for the other cell types [30], which was also used in our simulations. The developers of bMIND also proposed an alternative testing procedure, in which they only detected the top cell type with the minimal differential expressed (DE) p-value within a gene. This testing procedure was supported by some single-nucleus RNA-sequencing (snRNA-Seq) studies [31, 32] which reported that most CTS DE genes are only differentially expressed in one single cell type. In contrast, our meQTL analyses aimed to capture not only meQTLs that are specific in one single cell type but also meQTLs that are shared across multiple or all cell types. Previous studies have reported the existence of a substantial proportion of meQTLs that exhibit this shared pattern across diverse cell types [1, 33]. Therefore, in our simulations we did not adopt the alternative testing procedure proposed by bMIND. Instead, the marginal conditional test model was fitted for TCA and bMIND to control FDR and to provide a fair comparison with HBI. Although HBI generally outperformed other methods in our QTL-based simulations, we note that the deconvolution step in TCA and bMIND can output sample-level CTS profiles which enables other sample-level analyses (e.g., CTS co-expression networks), while methods based on the interaction model, like HBI, do not have this additional output.

In real data applications, the use of stringent statistical thresholds and independent replication datasets [1] enables the identification of CTS-meQTLs with high confidence and generalizability. Specifically, our identified meQTLs were supported by high replication rates in isolated CD4+ T-cells, CD8+ T-cells, and monocytes. We highlighted one example of the potential of our approach to identify biologically relevant cell types for complex traits. The colocalization analyses between meQTLs and GWASs highlight SNP rs2395178 in *HLA-DRA*, which was identified as a CD8+ T-cell specific meQTL for cg00886432. *HLA-DRA* belongs to the human leukocyte antigen (HLA) complex family, which plays an important role in antigen presentation and immune defense [34], and is well known as the major genetic determinant of T1D [26]. Our colocalization analyses revealed that rs2395178 was shared between methylation at cg00886432 in CD8+ T-cells and T1D (PPFC=0.9802). There has also been evidence for the contribution of CD8+ T-cells to the progression of T1D by the destruction of pancreatic beta cells [28]. Altogether, our downstream analyses helped to explain the relationship between this SNP-CpG locus and T1D, especially in CD8+ T-cells. Similarly, for other complex traits, we also identified biologically relevant cell types through meQTL-GWAS colocalization, and our findings strongly agreed with heritability enrichment analyses [27].

We acknowledge several limitations of our study. First, similar to other methods [10-12], our model depends on cell type proportions *W*_*k*_ as an input, but currently, only the estimated proportions are used directly to approximate the biological ground truth, which introduces additional error. Although we showed in simulations that the performance of HBI was generally robust in the noisy settings (noised *W*_*k*_ was provided), incorporating uncertainty specifically for estimated cell type proportion is warranted in the future. Second, we used an meQTL dataset obtained from experimentally isolated white blood cells [1] as the “gold standard” to replicate our findings. However, their CpG data was generated using the Illumina Human Methylation 450K BeadChip while our results were based on the Illumina Infinium Methylation EPIC BeadChip. Also, only a total of 11.2 million SNP-CpG pairs that were preselected in their bulk meQTL analysis were available. As a result, not all our significant results were represented in their database, limiting our replication analysis to the subset of pairs that overlapped. Third, in colocalization enrichment analysis, we did not observe significant results for some traits (i.e., no significant enrichments for heart attack or stroke). The potential reasons might be that our CTS-meQTLs were only derived from white blood cells and may not cover the causal cell types, and the small number of identified colocalizations in some traits may impact our results as well. Therefore, re-applying HBI on a dataset with a larger sample size and a wider range of causal cell types will help to obtain a more powered and complete CTS-meQTL catalog [33].

## Methods

### Statistical model

We model the relationship between methylation level at one CpG and one SNP as:

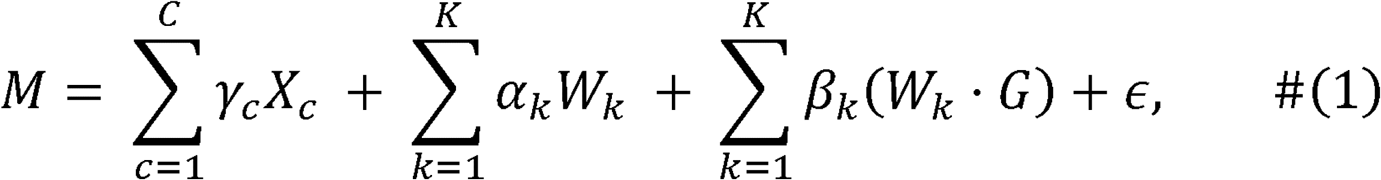

Where *M* is the bulk methylation level, *W*_*k*_ is the proportion of the *k* th cell type, *G* is the genotype of the SNP (the number of alternative alleles) of interest, *X*_*c*_ represents the *c*th covariate, such as age, sex, or ancestry, *ϵ* is a normally distributed error, and *α*_*k*_, *β*_*k*_,*γ*_*k*_ are regression coefficients. The coefficients of the interaction terms *β*_*k*_ are of primary interest: intuitively, if there exist genetic effects of DNAm in cell type *k*, the observed association between methylation and genotype should be stronger in samples with a higher fraction of cell type *k* compared to samples with lower fractions [12]. Of note, this model without intercept is equivalent to the following one, due to the fact that cell type proportions add up to 1:

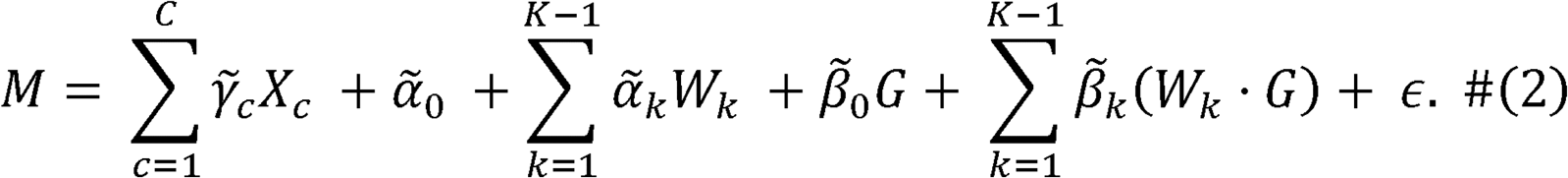

The difference between the two models lies in the interpretation of coefficients. In model (1) *β*_*k*_ represents the genetic effects on DNAm in cell type *k* (*k =* 1,2,…, *K)*. In model (2). 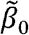, represents the genetic effects on DNAm in cell type *K* and 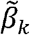 represents the changes in genetic effects in cell type *k* (*k =* 1,2,…, *K* −1)compared to the effect 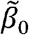 in cell type *K*. Therefore, 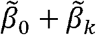 corresponds to the genetic effects in cell type *k* (*k =* 1,2,…, *K* −1). For simplicity, we use model (1) in the following derivations.

In order to take the sparsity of genetic effects into consideration and to update information derived from CTS methylation data from a small group of samples, a hierarchical framework is used to construct priors for regression coefficients [35]. To achieve optimal performance, we recommend that the small group of samples with CTS methylation data can be a subset of the overall samples with bulk data, or be similar samples drawn from the same study or cohort as the bulk samples. We first assume a multivariate normal distribution for coefficients for interaction terms ***β***:

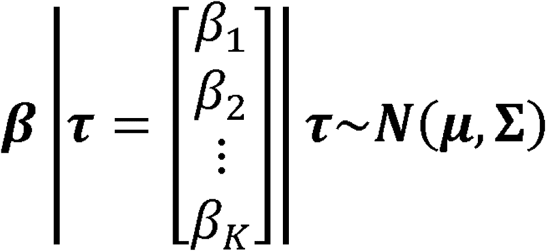

- **with general prior (no CTS methylation data)**

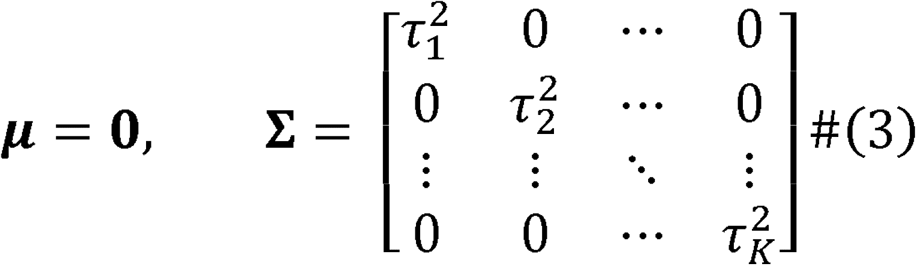

- **with prior derived from CTS methylation data of a small group of samples**

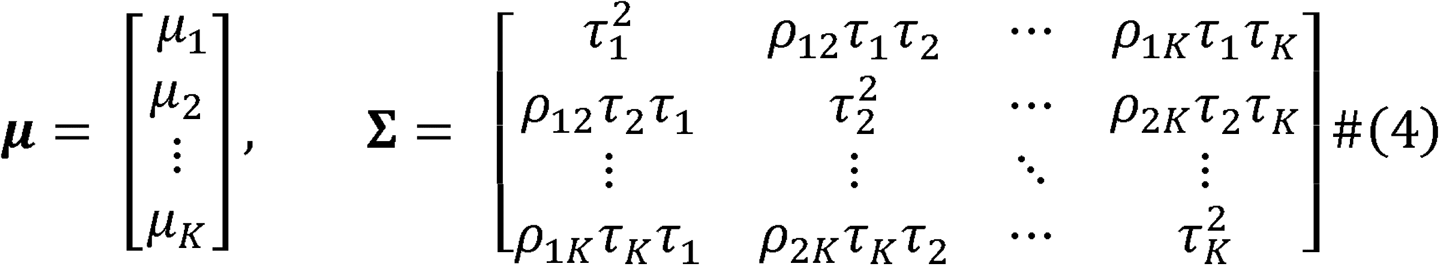

Where *µ*_*k*_ and *ρ*_*jk*_ are updated from the CTS methylation data in the following ways

(1) 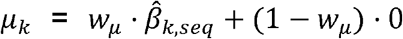, where 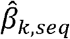 is the estimated effect when regressing the cell type *k* methylome on the SNP, and *w*_µ_ = 1 −*p*_adjust_ where *p*_adjust_ is the p-value adjusted by Benjamini & Hochberg (BH) or Bonferroni [17], as defined by users.

(2) 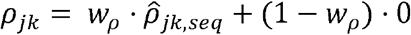, where 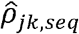 is the estimated genetic correlation between cell type *k* methylation and cell type *j* methylation, and *w*_*p*_= 1−*p*_*adjust*_ where *p*_*adjust*_ is the corresponding p-value for 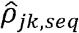, adjusted by Benjamini & Hochberg (BH) or Bonferroni [17], as defined by users.

In both settings with and without CTS methylomes derived from CTS DNAm data, the variable-specific parameter 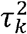 controls the degree of shrinkage: as 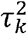 gets close to 0, *β*_*k*_ is shrunk to *µ*_*k*_, while as 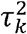 gets larger, the amount of shrinkage will be smaller. We further model 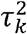 using the exponential distribution with variable-specific hyperparameters *s*_*k*_:

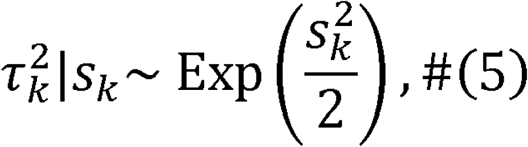

where *s*_*k*_ was modelled using a gamma distribution as a hyper-prior:

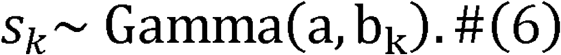

In this way, we allow different degrees of shrinkage for different variables by introducing the variable-specific parameters *s*_*k*_ and 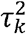. We also derive the conditional posterior distributions of *s*_*k*_ and 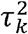 as follows:

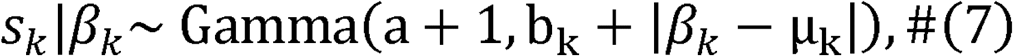

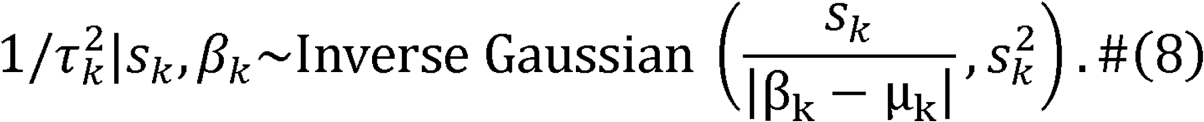

These will be used in the model fitting algorithm.

### EM-IWLS algorithm for model fitting and inference

We fit the hierarchical Bayesian interaction model by a modified iterative weighted least squares (IWLS) algorithm, proposed by Yi et al. [35]. Compared with usual IWLS, the new method incorporates an expectation–maximization (EM) algorithm that treats the unknown variances and 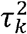 the hyperparameter *s*_*k*_ as missing data and estimates the *β* by averaging over these missing values, hence it is also referred to as the EM-IWLS algorithm.

In each iteration of the E-step, we update the missing values 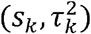 by their conditional expectations derived from (7) and (8). In the M-step, we update *β* by maximizing the expected log-likelihood. We need to incorporate the prior *β* | *τ* into the normal likelihood as additional data points [36]. Let *J* denote the total number of variables: (*J −K)* covariates (e.g., *α*_*k*_, *γ*_*c*_) included to address potential confounding, and *K* covariates (*β*)of our interest, and let ***θ* = [*γ***^***T***^, ***α***^***T***^, ***β***^***T***^]^***T***^ *∈****R***^***J***^. Model (1) could be expressed as:

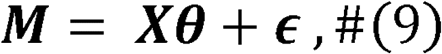

Where *X* ∈ ***R*** ^*n×J*^ is the original design matrix in model (1).

Then we update the regression coefficients by running the augmented linear regression:

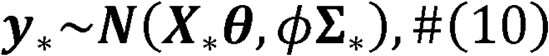

Where *y*_*\**_ = [***M***^***T***^,**0**^**T**^, **µ**^**T**^]^**T**^ is an ((*n* + *J*) ×1)vector of methylation levels for *n* samples and prior means for *J* covariates, 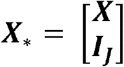 is an ((*n* + *J*) ×*J*) matrix constructed by the design matrix ***X*** in (9) and the identity matrix, and 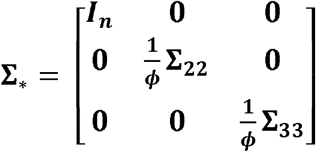 is an

((*n* + *J*) ×(*n* + *J)*) matrix with diag 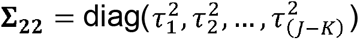 and Σ33 is the (*K* ×) prior variance matrix for *β* | *τ*. Then in each iteration, we can update the estimates:

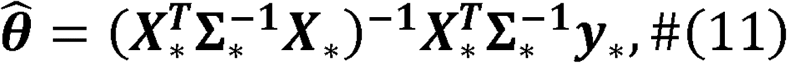

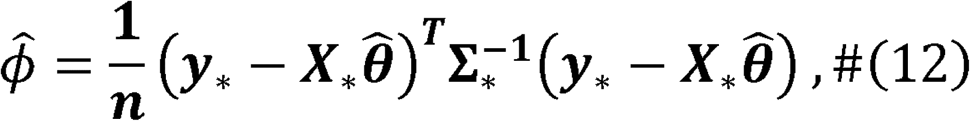

until convergence. We can also get the variance of regression coefficients:

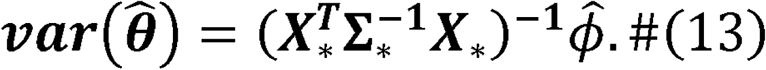

The EM-IWLS algorithm is summarized as follows.

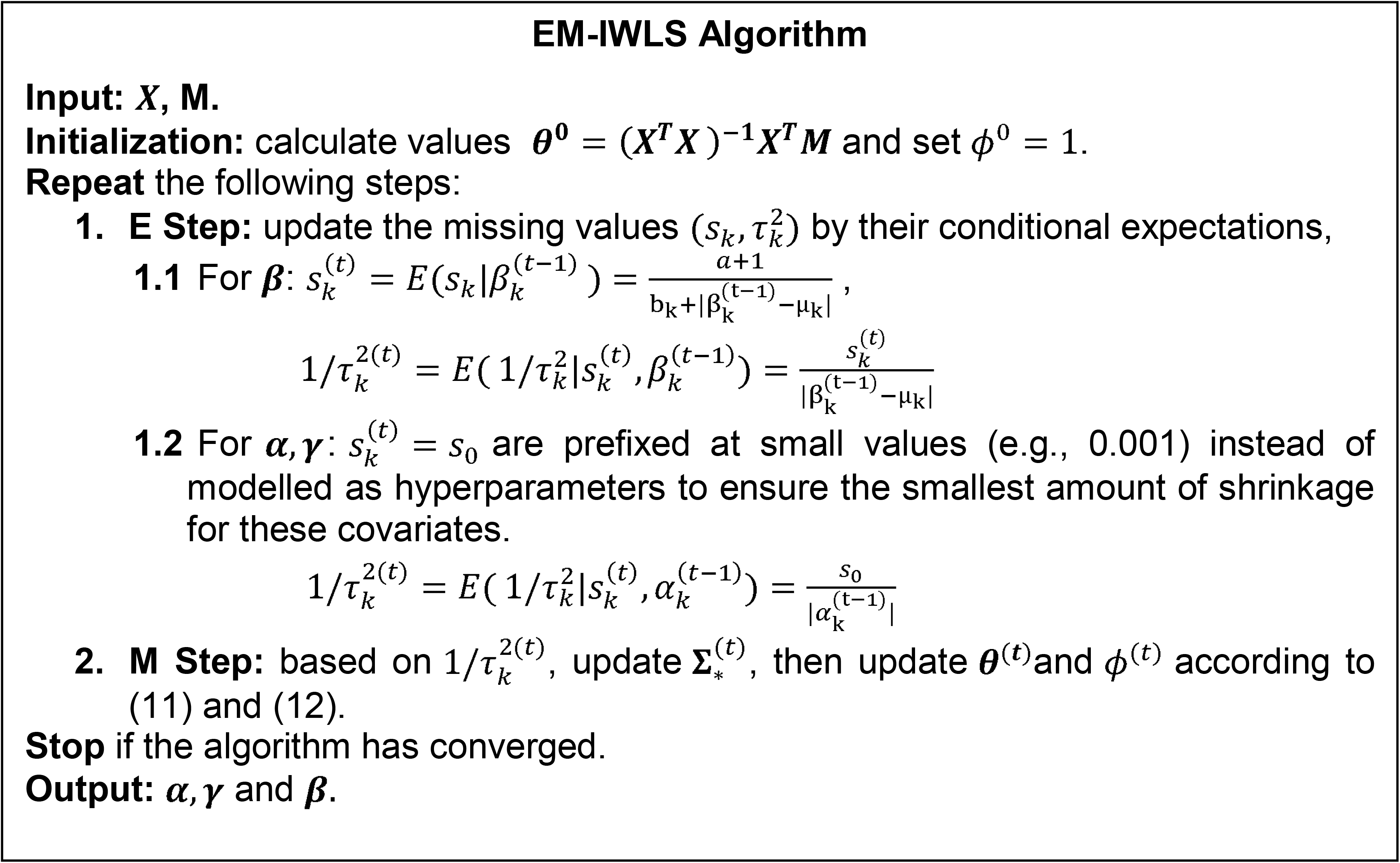

We define convergence as each element of |*θ*^(t)^ − *θ* ^(t −1)^|smaller than *δ*, with *δ*, to be a small value (e.g., 1E-05). 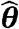 and 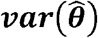 can then be obtained from the last updates.

For the choice of (a,b_*k*_), we fix a = 0.5 as the default since the overall degree of shrinkage can be determined by b_k_ [35]. For the user-defined b_k_, we suggest taking the moderate sample size (e.g., n = *0* (10^2^)), we suggest b_k_ = 0.2 for most cell types sample size and the abundance of the corresponding cell type into consideration. For (average of cell type proportions>10%), and b_k_ = 5 to induce a less informative prior for the least abundant cell type (average of cell type proportions<5%). Otherwise, the estimation of the coefficient for the least abundant cell type might be overwhelmingly driven by the prior. For larger sample sizes (e.g.,n = *O* (10^4^)) or much more abundant cell type, b_k_ could be decreased accordingly.

### Simulation settings

In this section, we introduce the simulation procedure to evaluate the performance of HBI. CTS DNAm in our simulations were generated based on genotype data from the Wellcome Trust Case Control Consortium (WTCCC) (n=15,918) [37]. In the cell type with genetic effects, the heritability of the DNAm was fixed as 0.3, the effect sizes of the causal SNPs were generated by a multivariate normal distribution [38], and GCTA [39] was applied to simulate the DNAm in this cell type. We also generated cell type proportions using a Dirichlet distribution for three cell types with parameters 5.30, 1.27, and 1.62. These parameters were chosen based on the suggestion from Li et al. that the mean cell type composition standard deviation is around 0.13, which was estimated from the Cibersort blood true proportions [40]. Then, for each sample, the bulk DNAm levels were computed as a weighted sum of the simulated CTS DNAm levels, weighted by the corresponding cell type proportions, plus an independent and identically distributed (iid) noise term *∈ ∼ N(0,0*.*01)*.

We considered three main scenarios, and in each of them, we assumed that the total number of SNPs near the simulated CpG site to be 500 and varied the proportion of causal SNPs from 10% to 20% to 40%. Each simulation setting was repeated 10 times.

(1) Scenario 1: there were genetic effects only in the major/most abundant cell type.

(2) Scenario 2: there were genetic effects only in the minor/least abundant cell type.

(3) Scenario 3: there were correlated genetic effects in all three cell types, and the genetic correlation among the cell types was set to 0.5.

We compared our method HBI with TCA [10], bMIND [11] and the basic interaction model, which fits model (1) directly using OLS. For HBI and the basic interaction model, we inputted the simulated bulk DNAm and cell type proportions and directly obtained the genetic effects for each cell type as the estimated coefficients for the interaction terms (*W*_*k*_ .*G*) For TCA and bMIND, we first inputted the bulk DNAm and cell type proportions to get deconvoluted CTS DNAm, and then tested the association between CTS DNAm and genotype using PLINK [41] to fit the following two models:

1. Marginal model which regresses the deconvoluted DNAm for cell *j*, 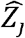, on the genotype *G* (equivalent to *marginal test* in TCA) [30]:

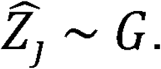

2. Conditional model which regresses the deconvoluted DNAm for cell *j* on the genotype with DNAm for all other cell types controlled (equivalent to *marginal conditional test* in TCA):

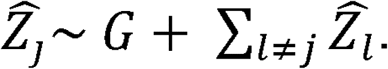

The coefficients for genotype *G* would then be the estimated genetic effects in cell *j* .CTS-meQTLs were identified with FDR controlled at 0.05 for each cell type. In each setting, the performance of different methods was compared in terms of correlation between the estimated and true effect sizes, the MSE between the estimated and true effect sizes, power, and FDR calculated as follows:

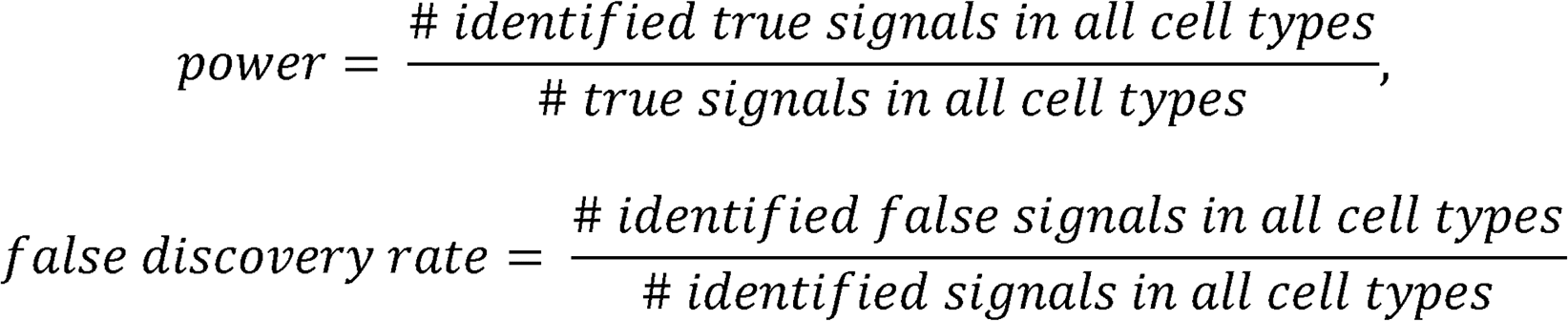

Both HBI and bMIND had the optional step to incorporate CTS information to update priors (derived from cell-sorted MC-seq data in our case and from scRNA-seq data in bMIND’s original case). Here we also assumed that for a small proportion (5%) of all samples, their CTS methylation data were available. In each simulation setting, we further compared HBI and bMIND both without this information incorporated and with this information incorporated (denoted as HBI_CTS-prior, bMIND_CTS-prior).

Since all methods included here relied on cell type proportions, we further evaluated the robustness of all methods when noisy cell type proportions were given. With the proportion of causal SNPs fixed as 20% in scenario 3, we randomly simulated noise from a left-truncated normal distribution (truncation point is zero), added noise to the true cell type proportions, and then normalized the sum of proportions to be 1. Two additional simulation settings were performed as we adjusted the standard deviation of the added noise so that the generated noisy cell type proportions would have mean absolute error (MAE) of 0.05 and 0.1, respectively.

### Study cohort for real data applications

The Women’s Interagency HIV Study (WIHS) is a multi-center, prospective, observational cohort study [15]. All participants are women with HIV or at risk for HIV acquisition. Informed consent was provided by all WIHS participants via protocols approved by institutional review committees at each affiliated institution. In our analysis, participants with matched genetic data and bulk DNA methylation measured in PBMC (n=431) and a separate group of participants with CTS DNA methylation data (n=47) were included. Demographic and clinical characteristics are summarized in **Supplementary Table 1**.

### Genotyping, imputation, and quality control

The WIHS sample were genotyped using the Infinium Omni2.5 Bead-Chip that targeted approximately 2.4 million SNPs. Minimac4 was used for imputation with the 1000 Genomes Project 3 as the reference panel [42, 43]. We removed SNPs with minor allele frequency<0.05, missing rate>5%, imputation quality r^2^<0.8, and those that deviated significantly from Hardy–Weinberg equilibrium (p<1e–6). As a result, approximately 4.6 million SNPs passed QC and were used for CTS-meQTL estimation.

### DNA methylation

DNA methylation measured using DNA isolated from PBMC was profiled using the Illumina Infinium MethylationEPIC BeadChip. We followed methods described in Lehne et al. [44] to perform methylation normalization and adjust for potential batch effects. A total of 852,073 CpGs for the 431 individuals passed quality control steps and were used as bulk DNAm data. We applied the method described by Houseman et al. to estimate the cell type proportions for CD4+ T-cells, CD8+ T-cells, natural killer cells, B cells, monocytes, and granulocytes [19, 45]. Another separate group of the WIHS cohort (n=47) were isolated for CD4+ T-cells, CD8+ T-cells, and monocytes. DNAm for each sorted cell type was profiled by the Agilent SureSelectXT Methyl-seq. After quality control and extracting CpGs that overlapped on both platforms, we had 390,851 CpGs measured in CD4+ T-cells (n=28), 385,679 CpGs measured in CD8+ T-cells (n=28), and 407,646 CpGs measured in monocytes (n=27), which were used as CTS DNAm data to update priors.

### CTS-meQTL estimation and replication

We applied HBI to identify CTS meQTLs in the WIHS cohort for six cell types: CD4+ T-cells, CD8+ T-cells, natural killer cells, B cells, monocytes, and granulocytes. For each CpG, we considered the following model for SNPs from 500kb upstream to 500kb downstream:

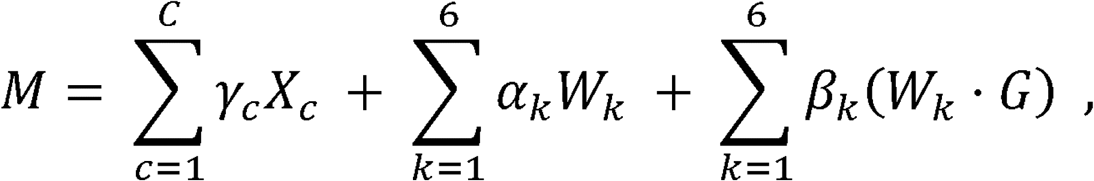

Where *M* is the bulk methylation M-value, *W*_*k*_ is the cell type proportion of the *k* th cell type, *G* is the genotype of the SNP, *X*_*c*_ is a collection of previously identified relevant covariates: age, estimated global ancestry, local ancestry [46], tobacco use, alcohol consumption, HIV infection status, log_10_ of HIV RNA viral load, the top 5 genotype principal components (PCs), and the top 10 PCs on DNA methylation levels of control probe. HBI was applied to estimate the regression coefficients in the above model, and for CD4+ T-cells, CD8+ T-cells, and monocytes, we further incorporated the priors derived from the CTS methylation data available in a small group of subjects. The choices of the parameters in the hyper prior *Gamma(a,b*_*k*_) were as follows: a = 0.5 for all cell types, b_k_ = 5 for granulocytes, b_k_ = 0.2 for natural killer cells, B cells, monocytes, and b_k_ = 0.05 for CD4+ T-cells, CD8+ T-cells. Among the 852,073 CpGs, a total of 1.4 billion SNP-CpG pairs were tested, and significant meQTLs were selected using Bonferroni correction (p<0.05/1,384,706,562/6=6.02E-12).

Independent data were used to replicate our identified CTS-meQTLs. We downloaded datasets for meQTLs in isolated white blood cell subsets (i.e., CD4+ T-cells, CD8+ T-cells, monocytes, neutrophils) (n=60 individuals) [1]. For our identified SNP-CpG pairs in the respective cell types, we calculated the percentage of pairs that were significant in the replication set (p<0.05), the percentage of pairs with directional consistency in effect sizes, and the percentage of replicated pairs (p<0.05 and same effect direction). Among the replicated pairs, we also calculated the correlations of the effect sizes.

### MeQTL enrichment in genomic functional annotations

For all the variants tested for SNP-CpG associations, we used annotatr [47] and its built-in annotation databases to make CpG annotations (CGI, CGI shelves, CGI shores, inter CGI regions), gene body annotations (regions<1kb upstream of the transcription start site, coding sequence, exons, introns, intergenic regions, 5’UTRs, 3’UTRs), gene regulatory and open chromatin annotations (active promotor, weak promotor, strong enhancer, weak enhancer, insulator, regions with heterochromatic or heterochromatin-like characteristics). For gene regulatory and open chromatin annotations [48], we used the database for the K562 cell line, which is commonly used to study hematopoiesis [49]. To test whether the identified meQTLs were enriched in some functional regions, we performed functional enrichment analysis using Fisher’s exact test [50, 51]. A 2×2 contingency table was built as follows:

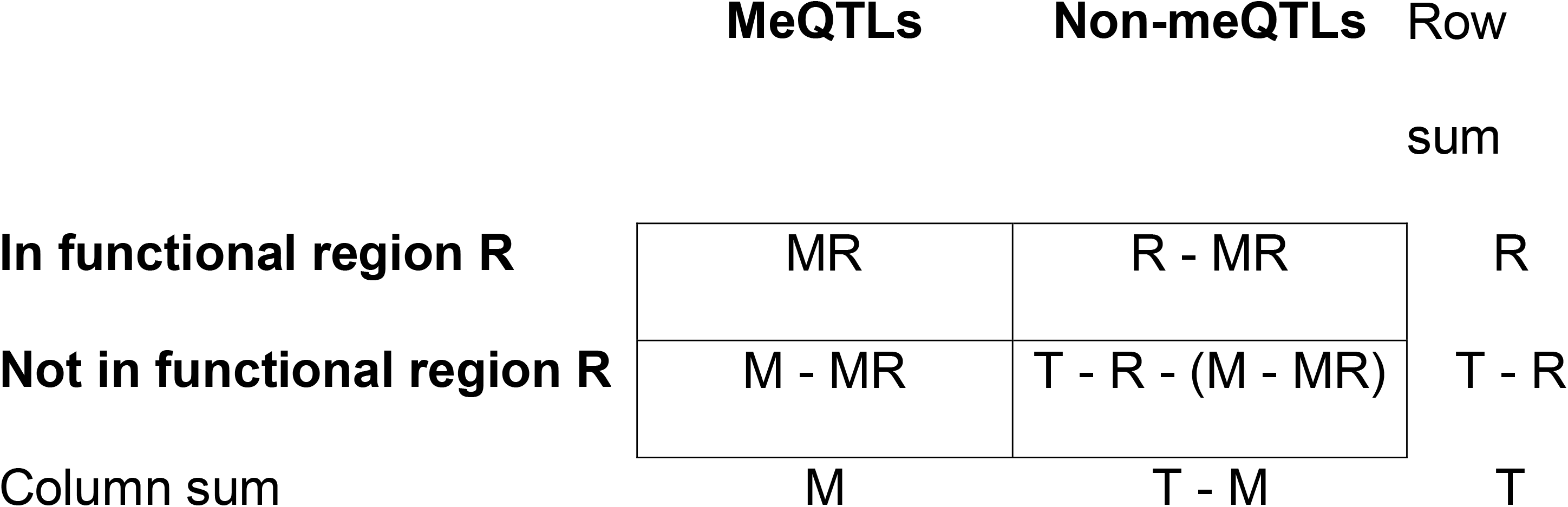

The total sum of the contingency table (T) was the number of variants that were tested for SNP-CpG associations. The number of identified meQTLs that were mapped in one specific functional region corresponds to the upper-left cell of the table (MR). The remaining three cells of the table can be calculated based on MR and the row/column sums. Based on the 2×2 contingency table, we tested whether the meQTLs were enriched in the functional region more often than by chance expected by the genome background (non-meQTLs) [52, 53]. For each cell type, we performed this analysis separately and derived the enrichment estimates as log of odds ratios and its 95% confidence intervals. Enrichment across all cell types was conducted by combining CTS-meQTLs into a union set comprising meQTLs identified in at least one cell type.

### Pathway analyses based on identified CTS-meQTLs

For the identified meQTLs in each cell type, we used ANNOVAR to map variants to their nearest gene, and for variants in intergenic regions, the closest gene was kept [54]. Pathway enrichment analyses were conducted with QIAGEN Ingenuity Pathway Analysis (IPA) (QIAGEN Inc., https://digitalinsights.qiagen.com/IPA) [21]. In each cell type, we reported significant pathways at Bonferroni-adjusted p<0.05.

### Colocalization of meQTL with GWAS loci

To identify potential associations between meQTLs and complex traits, we applied HyPrColoc (Hypothesis Prioritization for multi-trait Colocalization) [24] in multiple genomic regions. We downloaded GWAS summary statistics published by Barbeira et al. [25, 55], and used the 57 traits in the categories of blood cell counts, cardiometabolic, immune, and allergy. Since colocalization reports the posterior probability that two traits are colocalized in a specific linkage disequilibrium (LD) region [24, 56], we first performed clumping on the meQTLs identified in each cell type. For each CpG, highly correlated genetic variants were clustered into one clump with an LD r^2^>0.1 [57], resulting in 7,766 meQTL clumps for CD4+ T-cells, 4,211 clumps for CD8+ T-cells, 4,568 clumps for monocytes, 3,219 clumps for B cells, 2,649 clumps for natural killer cells, and 1,821 clumps for granulocytes. For each cell type, the genetic variants in each meQTL clump were matched with GWAS summary statistics. Then for each meQTL-GWAS region pair, HyPrColoc was applied on the effect size and the corresponding standard errors. The PPFC was used to identify significant (PPFC>0.50) colocalizations [33, 58].

### Cell type specific enrichment in meQTL-GWAS colocalizations

To investigate the cellular specificity of complex traits, we performed enrichment analyses to study if the meQTL-GWAS colocalizations for each trait were enriched in certain cell types. Here, meQTLs in granulocytes were excluded due to low numbers of colocalizations identified across traits, and meQTLs in bulk level (282,965 clumps) were included to assess if CTS-meQTLs can reveal more cellular specific information. We also excluded three traits with a very small number of meQTL-GWAS colocalizations (<10) across all cell types. As a result, 54 of the 57 GWAS traits remained in the enrichment analyses. For each trait, in each cell type the enrichment score was defined as the ratio between the percentage of meQTL-GWAS colocalizations (colocalized meQTL clumps) in that cell type and the percentage of meQTL clumps covered by that cell type:

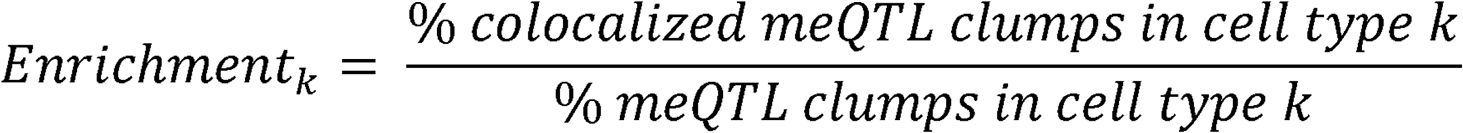

To determine significant colocalization enrichments in certain cell types, the test for equality of proportions with continuity correction was performed to test if *Enrichment*_*k*_ *> 1* (p< 0.05/54/6=1.5E-04). To evaluate the identified enrichment results, for the same GWAS traits we also performed heritability enrichment analyses using 66 functional annotations from GenoSkyline-Plus (v1.0.0)[27]. For our identified traits with colocalizations enriched in certain cell types, we determined if the heritability of this trait also enriched in this cell type at p<0.05.

## Declarations

### Ethics approval and consent to participate

The study was approved by the committee of Human Research Subject Protection at Yale University and the Institutional Research Board Committee of the Connecticut Veteran Healthcare System. Informed consent was provided by all WIHS participants via protocols approved by institutional review committees at each affiliated institution.

### Consent for publication

*Not applicable*

### Availability of data and materials

Genotype data used in the simulation were downloaded from the Wellcome Trust Case-Control Consortium (https://www.wtccc.org.uk). Data in the application part were from the WIHS cohort, which has been identified as one with multiple vulnerabilities (e.g., racial/ethnic minority women, coinfected). Whereas participants from the cohort who contributed to the findings summarized in this manuscript provided written consent for genetic studies, said consent was collected prior to the most recent guidelines and requirements for data sharing. The WIHS cohort operates under an alternative data sharing plan registered with the National Institutes of Health and access to data can be requested by submitting a Concept Sheet, which can be found along with instructions for Concept Sheet submission, at https://statepi.jhsph.edu/mwccs/. The accession number for the WIHS in dbGaP genomic data is now provided (phs001503). The cohort is currently being re-approached to obtain informed consent for sharing of their data. This has been consistent with other genomic studies in the WIHS cohort. The algorithm for HBI and codes for analysis are on GitHub.

## Authors’ contributions

YC, BC and HL developed the statistical framework. YC implemented the algorithm and performed statistical analysis. HL and XZ assisted in data analysis. BEA, GD, SS, AE, JM, MF, SK, KA, and MC provided DNA samples and contributed to manuscript preparation. KX and HZ were responsible for the study design. KX advised on sample preparation and the biological interpretation of findings. HZ advised on statistical and genetics issues. YC drafted the manuscript. All authors contributed to manuscript editing and approved the manuscript.

## Acknowledgements

The authors appreciate the support of the WIHS cohort and Yale Center of Genomic Analysis. Data in the application part of this manuscript were collected by the Women’s Interagency HIV Study (WIHS), now the MACS/WIHS Combined Cohort Study (MWCCS). The contents of this publication are solely the responsibility of the authors and do not represent the official views of the National Institutes of Health (NIH). MWCCS (Principal Investigators): Atlanta CRS (Ighovwerha Ofotokun, Anandi Sheth, and Gina Wingood), U01-HL146241; Baltimore CRS (Todd Brown and Joseph Margolick), U01-HL146201; Bronx CRS (Kathryn Anastos, David Hanna, and Anjali Sharma), U01-HL146204; Brooklyn CRS (Deborah Gustafson and Tracey Wilson), U01-HL146202; Data Analysis and Coordination Center (Gypsyamber D’Souza, Stephen Gange and Elizabeth Topper), U01-HL146193; Chicago-Cook County CRS (Mardge Cohen and Audrey French), U01-HL146245; Chicago-Northwestern CRS (Steven Wolinsky, Frank Palella, and Valentina Stosor), U01-HL146240; Northern California CRS (Bradley Aouizerat, Jennifer Price, and Phyllis Tien), U01-HL146242; Los Angeles CRS (Roger Detels and Matthew Mimiaga), U01-HL146333; Metropolitan Washington CRS (Seble Kassaye and Daniel Merenstein), U01-HL146205; Miami CRS (Maria Alcaide, Margaret Fischl, and Deborah Jones), U01-HL146203; Pittsburgh CRS (Jeremy Martinson and Charles Rinaldo), U01-HL146208; UAB-MS CRS (Mirjam-Colette Kempf, Jodie Dionne-Odom, Deborah Konkle-Parker, and James B. Brock), U01-HL146192; UNC CRS (Adaora Adimora and Michelle Floris-Moore), U01-HL146194. The MWCCS is funded primarily by the National Heart, Lung, and Blood Institute (NHLBI), with additional co-funding from the Eunice Kennedy Shriver National Institute Of Child Health & Human Development (NICHD), National Institute On Aging (NIA), National Institute Of Dental & Craniofacial Research (NIDCR), National Institute Of Allergy And Infectious Diseases (NIAID), National Institute Of Neurological Disorders And Stroke (NINDS), National Institute Of Mental Health (NIMH), National Institute On Drug Abuse (NIDA), National Institute Of Nursing Research (NINR), National Cancer Institute (NCI), National Institute on Alcohol Abuse and Alcoholism (NIAAA), National Institute on Deafness and Other Communication Disorders (NIDCD), National Institute of Diabetes and Digestive and Kidney Diseases (NIDDK), National Institute on Minority Health and Health Disparities (NIMHD), and in coordination and alignment with the research priorities of the National Institutes of Health, Office of AIDS Research (OAR). MWCCS data collection is also supported by UL1-TR000004 (UCSF CTSA), UL1-TR003098 (JHU ICTR), UL1-TR001881 (UCLA CTSI), P30-AI-050409 (Atlanta CFAR), P30-AI-073961 (Miami CFAR), P30-AI-050410 (UNC CFAR), P30-AI-027767 (UAB CFAR), P30-MH-116867 (Miami CHARM), UL1-TR001409 (DC CTSA), KL2-TR001432 (DC CTSA), and TL1-TR001431 (DC CTSA).

The authors gratefully acknowledge the contributions of the study participants and dedication of the staff at the MWCCS sites.

