## Supplementary Figures for "A hierarchical Bayesian interaction model to estimate cell-type-specific methylation quantitative trait loci incorporating priors from cell-sorted bisulfite sequencing data"

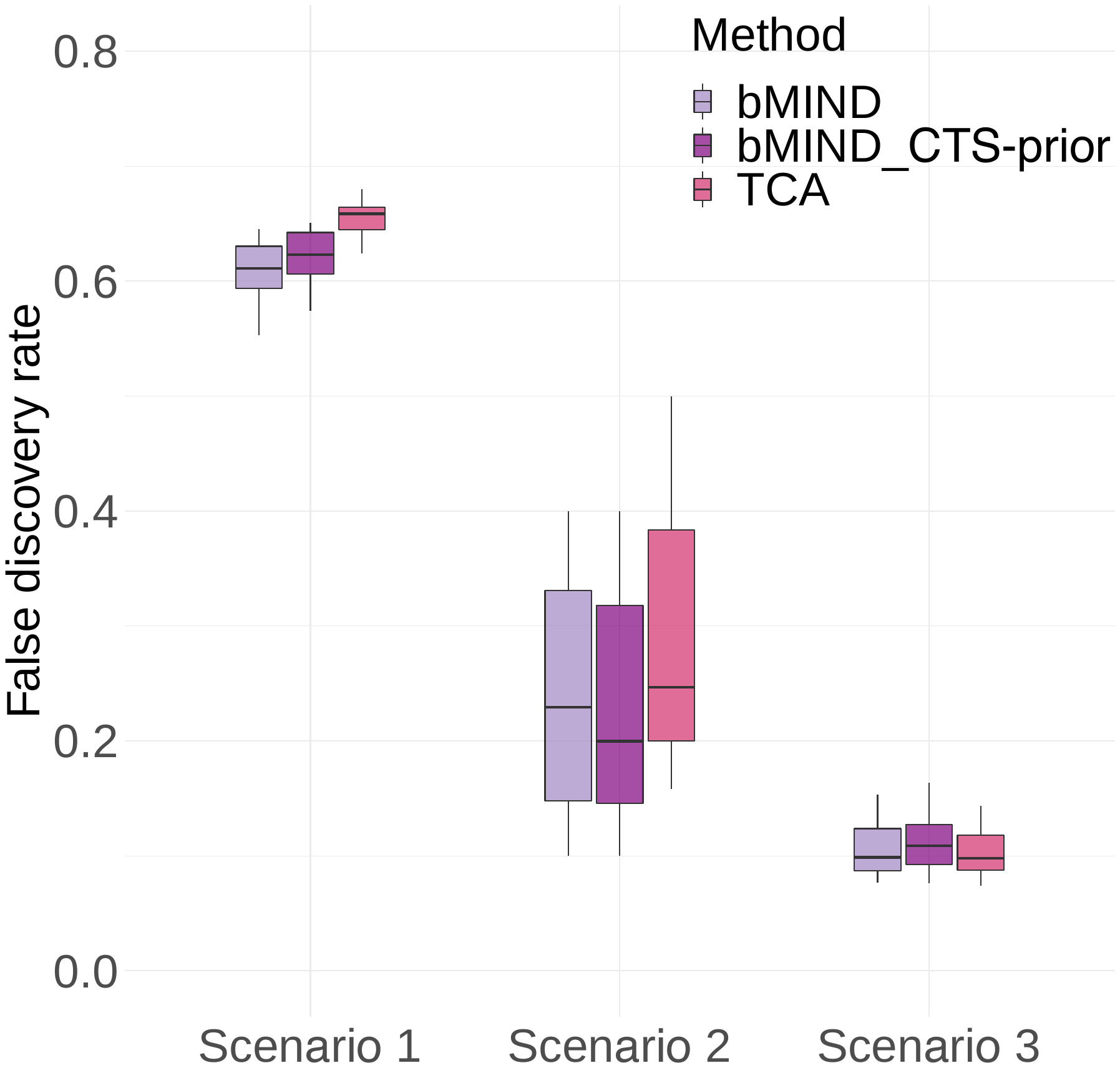

**Supplementary Figure 1: False discovery rate (FDR) of bMIND and TCA when using the marginal test.** Marginal test refers to the model that regresses the deconvoluted DNAm for one cell type on the genotypes, without controlling for all other cell types. FDR is presented in scenarios with genetic effects only in the most abundant cell type (Scenario 1), only in the least abundant cell type (Scenario 2), and with correlated genetic effects in all cell types (Scenario 3). In each scenario, the proportion of causal SNPs is 20%. bMIND_CTS-prior represents the version of bMIND with cell-type-specific (CTS) methylation data incorporated.

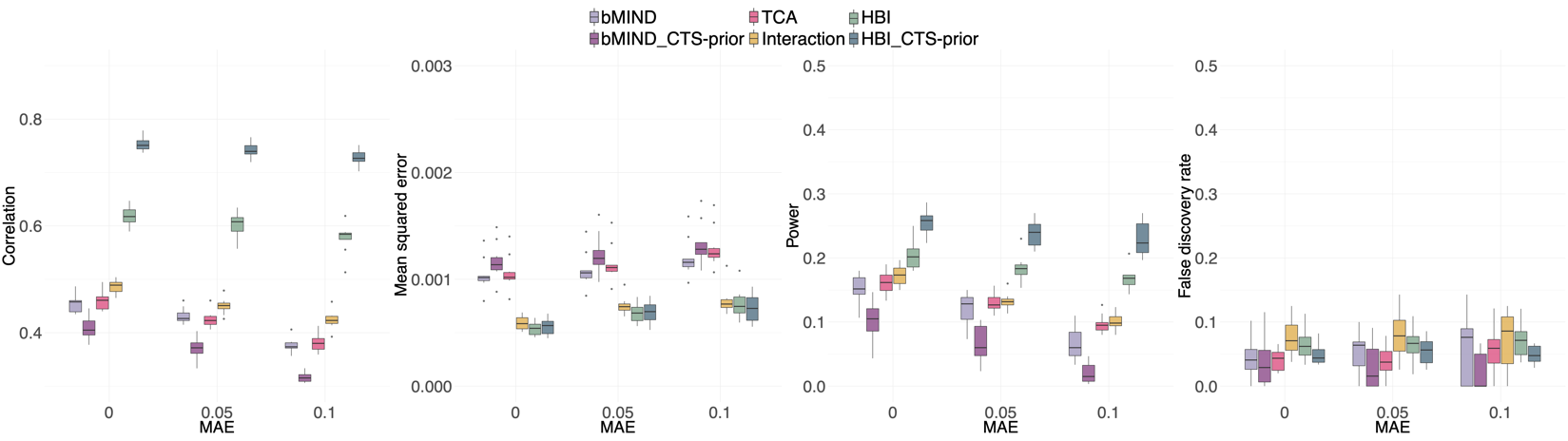

**Supplementary Figure 2: Performance in estimating cell type specific (CTS)-meQTLs with noisy cell type proportions.** We randomly simulate noise from a normal distribution, add noise to the true cell type proportions, and then normalize the sum of proportions to be 1. We adjust the standard deviation of the added noise so that the generated noisy cell type proportions would have mean absolute error (MAE) of 0.05 and 0.1. From left to right: correlation between estimated and true effect sizes, mean squared error (MSE) between estimated and true effect sizes, power, and false discovery rate as a function of MAE. Scenario 3 (correlated genetic effects in all cell types) is shown, and the proportion of causal SNPs is 20%. HBI_CTS-prior, bMIND_CTS-prior represent the version of the corresponding methods with CTS methylation data incorporated.

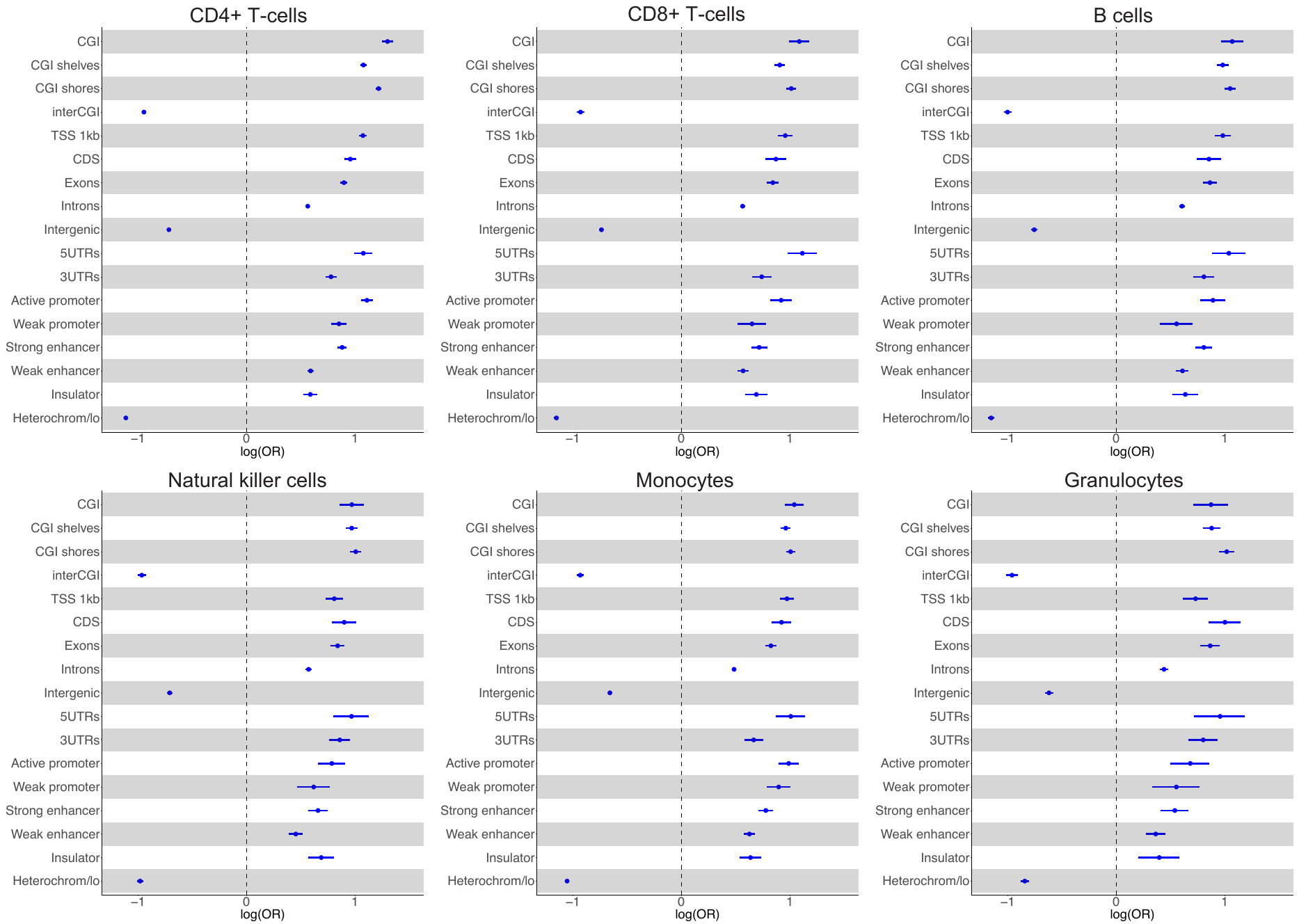

**Supplementary Figure 3: Functional enrichment for cell type specific (CTS)-meQTLs in CpG island (CGI) regions, gene body regions, and gene regulatory regions.** The logarithm of odds ratio (OR) with 95% confidence interval is presented. TSS 1kb: <1kb upstream of the transcription start site (TSS); CDS: coding sequence; UTR: untranslated exon region; Heterochrom/lo: regions that exhibit heterochromatic or heterochromatin-like characteristics.

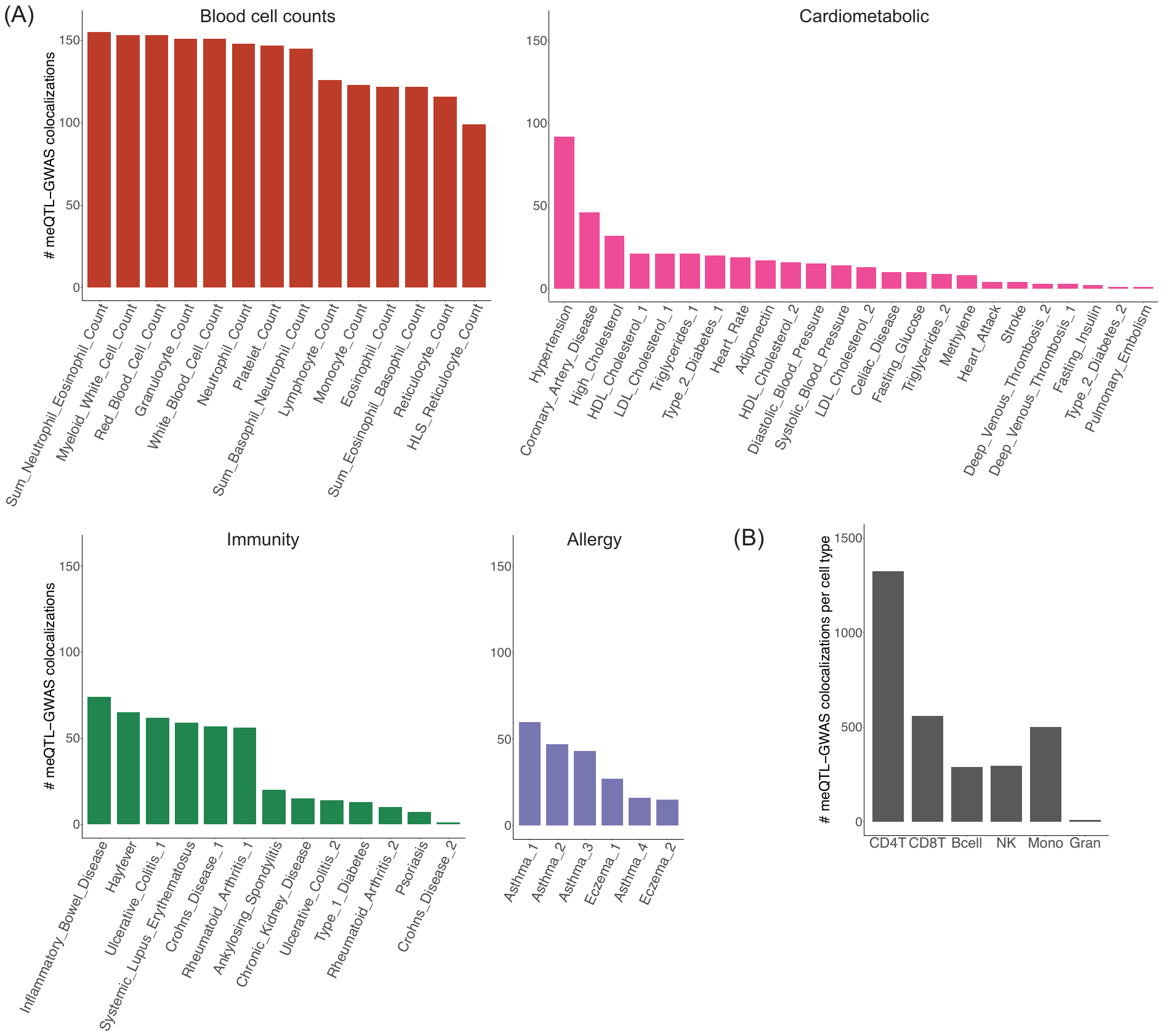

**Supplementary Figure 4: Colocalization of meQTLs with GWAS traits. (A)** Bar plot represents the number of meQTL-GWAS colocalizations per GWAS trait across all six cell types. **(B)** Bar plot represents the number of meQTL-GWAS colocalizations per cell type across all GWAS traits.

HLS reticulocyte: high light scatter reticulocyte; HDL: high-density lipoprotein; LDL: low-density lipoprotein; GWAS: genome-wide association studies; CD4T: CD4+ T-cells; CD8T: CD8+ T-cells; NK: natural killer cells; Mono: monocytes; Gran: granulocytes.

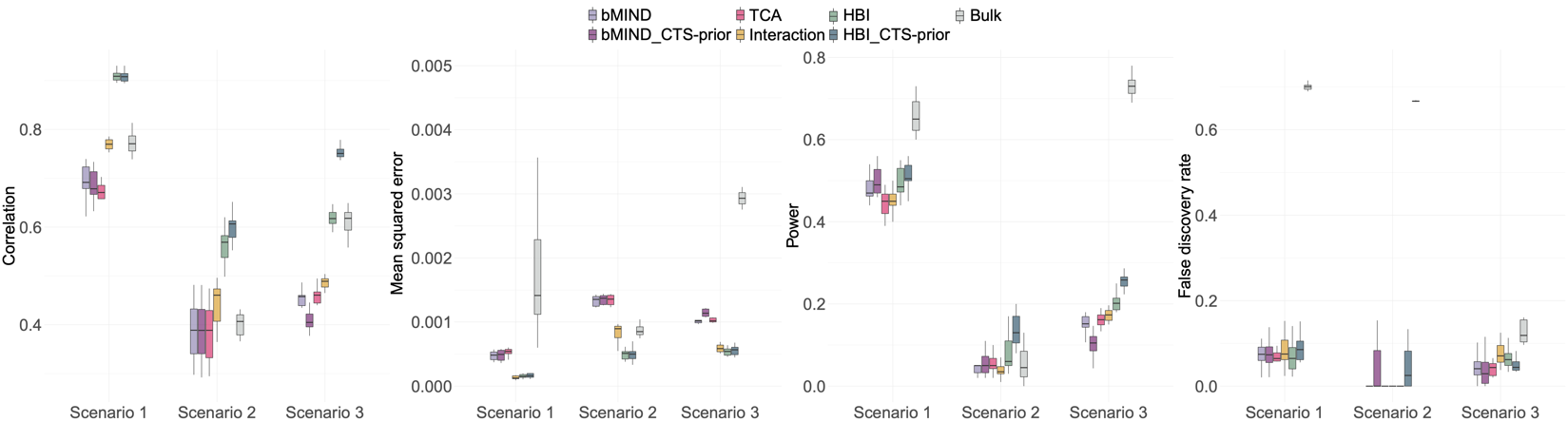

**Supplementary Figure 5: Comparison between cell type specific (CTS)-meQTLs and bulk meQTLs identified using the bulk data.** Bulk meQTLs are identified using the linear regression model that directly regresses bulk methylation on genotypes. To compare with CTS-meQTLs, we assume the identified bulk meQTLs are the same (homogeneous) in all cell types. Correlation, mean squared error (MSE) between estimated and true effect sizes, power, and false discovery rate are presented in scenarios with genetic effects only in the most abundant cell type (Scenario 1), only in the least abundant cell type (Scenario 2), and with correlated genetic effects in all cell types (Scenario 3). In each scenario, the proportion of causal SNPs is 20%. HBI_CTS-prior, bMIND_CTS-prior represent the version of the corresponding methods with CTS methylation data incorporated.
